# Patterns and drivers of the diving behavior of large pelagic predators

**DOI:** 10.1101/2022.12.27.521953

**Authors:** Amy Nuno, Jérome Guiet, Brooke Baranek, Daniele Bianchi

**Affiliations:** Department of Atmospheric and Oceanic Sciences, University of California Los Angeles, Los Angeles, CA, USA; Department of Earth System Science, University of California, Irvine, CA, USA

**Keywords:** Large marine predators, Diving depth, Vertical behavior, Physiology, Predator-prey interactions, Tuna, Shark, Billfish

## Abstract

Many large pelagic predators, including shark, tuna, and billfish, periodically dive to deep oceanic layers, connecting the surface and mesopelagic ecosystems. However, the patterns and drivers of deep dives across species remain poorly understood. To close this gap, we conduct a meta-analysis of the diving behavior of 24 pelagic predator species from the global ocean, resulting in 671 independent diving depth estimates from 87 tagging studies. Our analysis reveals consistent large-scale patterns in diving depths, with predators diving deeper offshore and during the day, and shallower closer to the coast and during the night. Deep diving species show similar diving depths during the night, with deeper dives for sharks, and shallower dives for tuna and swordfish. These patterns are reversed during the day, widening the gap between day and night vertical ranges for these groups. In contrast, shallow diving species show smaller variations between day and night dives, with sharks diving slightly deeper on average, followed by tuna and billfish. Correlations with co-located environmental variables suggest an important predictive role for proxies of prey abundance and light availability, as well as variables that influence physiology, such as oxygen and temperature. These relationships are more robust for deep divers during the day, and shallow divers at night. Our analysis highlights the value of tagging observations for the development of a mechanistic, quantitative characterization of vertical habitat use of large marine predators and its environmental constraints.

## Introduction

Large pelagic predators (LPP), including keystone species such as sharks, billfish and tuna, complete regular deep dives to depths of hundreds of meters below the surface (Andrzejaczek et al. 2022; Braun et al. 2022). These dives connect warm, well-lit epipelagic layers to the cold and dark mesopelagic ocean. By linking these distinct habitats via feeding at one depth and excretion at another, LPP influence biomass flows in marine food-webs, shape ecological interactions between species, and affect nutrients and carbon cycles (Doughty et al. 2016; Braun et al. 2022; Pinti et al. 2022).

Invisible to the human eye or even not known for a long time, the deep dives of LPP remained poorly characterized until the application of tagging approaches. From early technology based on active acoustics in the 70s and 80s, tags evolved to record the movement of marine animals in space and time, and today consist of sophisticated sensors recording variables that include pressure, temperature, and light. Tagging studies have revealed a broad spectrum of diving behaviors, ranging from sporadic dives of few minutes in duration, to diel dives following the diurnal cycle of solar illumination (Andrzejaczek et al. 2019).

The ultimate cause of these vertical excursions remains unclear, but is generally thought to involve a combination of feeding behavior (Musyl et al. 2003; Shepard et al. 2006; Vetter et al. 2008; Dewar et al. 2011; Williams et al. 2015; Francis et al. 2015a), energy conservation (Campana Steven E. and Dorey, 2011; Coffey et al., 2017; Hino et al., 2019; Kitagawa et al., 2004; Weng et al., 2009), orientation and navigation (Willis et al. 2009; Nasby-Lucas et al. 2009), and social interactions (Jorgensen et al. 2012).

Several proximate drivers of diving behavior have been suggested. For instance, temperature modulates physiological rates, including cardiac function and neural activity, and often determines the preferred depth range outside of which the survival of LPP may be compromised (Pinsky et al. 2020; Fredston et al. 2021). Because it decreases with depth, water temperature can limit or even prevent deep dives (Patterson et al. 2008; Musyl et al. 2011; Chiang et al. 2011; Hoolihan et al. 2015; Weng et al. 2017; Vaudo et al. 2018). Like temperature, oxygen decreases with depth over most of the ocean, constraining species physiology, ecology, and diving behavior (Brill 1994; Deutsch et al. 2020). The vertical diving patterns of LPP could thus be bounded by species-dependent oxygen tolerances, with shallow low-oxygen layers leading to shallow diving depths (Schaefer et al. 2009; Abascal et al. 2011). Beyond physiological constraints, prey occurrence is also an important driver of diving behavior, especially in relation to the distribution of mesopelagic fish that migrate to the surface at night (Musyl et al. 2003; Abecassis et al. 2012; Heard et al. 2018). Feeding on mesopelagic fish generally requires adaptations to detect prey in dimly lit environments (Abascal et al. 2010; Dewar et al. 2011). The physical characteristics of the environment such as fronts and mesoscale eddies also influence the occurrence of prey (Lévy et al. 2018) and thus the behavior of predators (Braun et al. 2019; Arostegui et al. 2022). While recent reviews summarize the significance of these drivers and their variability on time scales ranging from minutes to seasons (Andrzejaczek et al. 2019; Braun et al. 2022), systematic, quantitative assessments across species based on tagging data remain limited.

A mechanistic understanding of the drivers of LPP vertical migration could help in anticipating changes in the behavior of key species in a warming ocean (Hazen et al. 2019), in particular in the poorly observed mesopelagic ecosystem (Hidalgo and Browman 2019). LPP include closely monitored species of economic value for fisheries and recreation (Arostegui et al. 2022; Juan-Jordá et al. 2022), protected species, and species threatened by bycatch (Scales et al. 2018). Understanding their vertical behavior could thus help management of valuable stocks. A better representation of vertical habitat utilization and predator-prey interactions would also improve marine ecosystem models used to project the evolution of wild fisheries in a changing ocean (Lotze et al. 2019; Heneghan et al. 2021; Tittensor et al. 2021), and expand our understanding of the influence of LPP on biogeochemical cycles (Aumont et al. 2018; Bianchi et al. 2021; Pinti et al. 2022).

Here, we present a meta-analysis of existing tagging studies that identifies patterns and first-order drivers of the diving behavior of 24 pelagic predators belonging to shark, billfish, and tuna taxonomic groups. The analysis reveals distinct inter- and intra-specific sensitivities to environmental variables, which we discuss in relation to previously proposed hypotheses on the factors controlling deep diving behavior.

## Methods

### Meta-analysis overview

We conducted a meta-analysis of diving behavior from tagging studies based on a systematic literature review protocol (Haddaway et al., 2020). We selected relevant publications based on a list of keywords related to diving behavior of LPP. Based on a general analysis of the selected publications, we pre-determined the quantities to extract from each study, and the approaches required to extract them. These quantities consist of the preferred daytime and nighttime diving depths, total diving range, and matching spatial and temporal information. The extraction was replicated independently by three of the co-authors, and the data merged for analysis and comparison with co-located climatological environmental data.

We identified publications documenting LPP diving behavior by searching the Web of Science and Google Scholar with keywords that included “diel vertical migration”, “vertical behavior”, “tagging”, associated with common names for marine predator groups (e.g., tuna, shark, billfish, see Supplementary Table S1). A total of 135 peer-reviewed publications were identified. From these, we further analyzed 87 papers that reported quantifiable measures of diving depths, together with the location and time of the tagging events (see Supplementary Table S2 for the complete list of papers). We focused on species classified as pelagic according to FishBase (Froese and Pauly 2022).

### Extraction of large pelagic predator diving depths

We found significant diversity in the type of diving information reported in the literature, since different studies focus on various aspects of animal behavior, from diel vertical migrations to horizontal migrations, feeding habits, and habitat use. The format of this data is also variable. In some instances, diving depths are directly reported in the text or in tables. More often, diving depths are shown by histograms of the relative occurrence of individuals at specific depths (see example in Fig.1a), or by explicit trajectories of the individual’s depth as a function of time (see example in Fig.1b).

**Figure 1.**
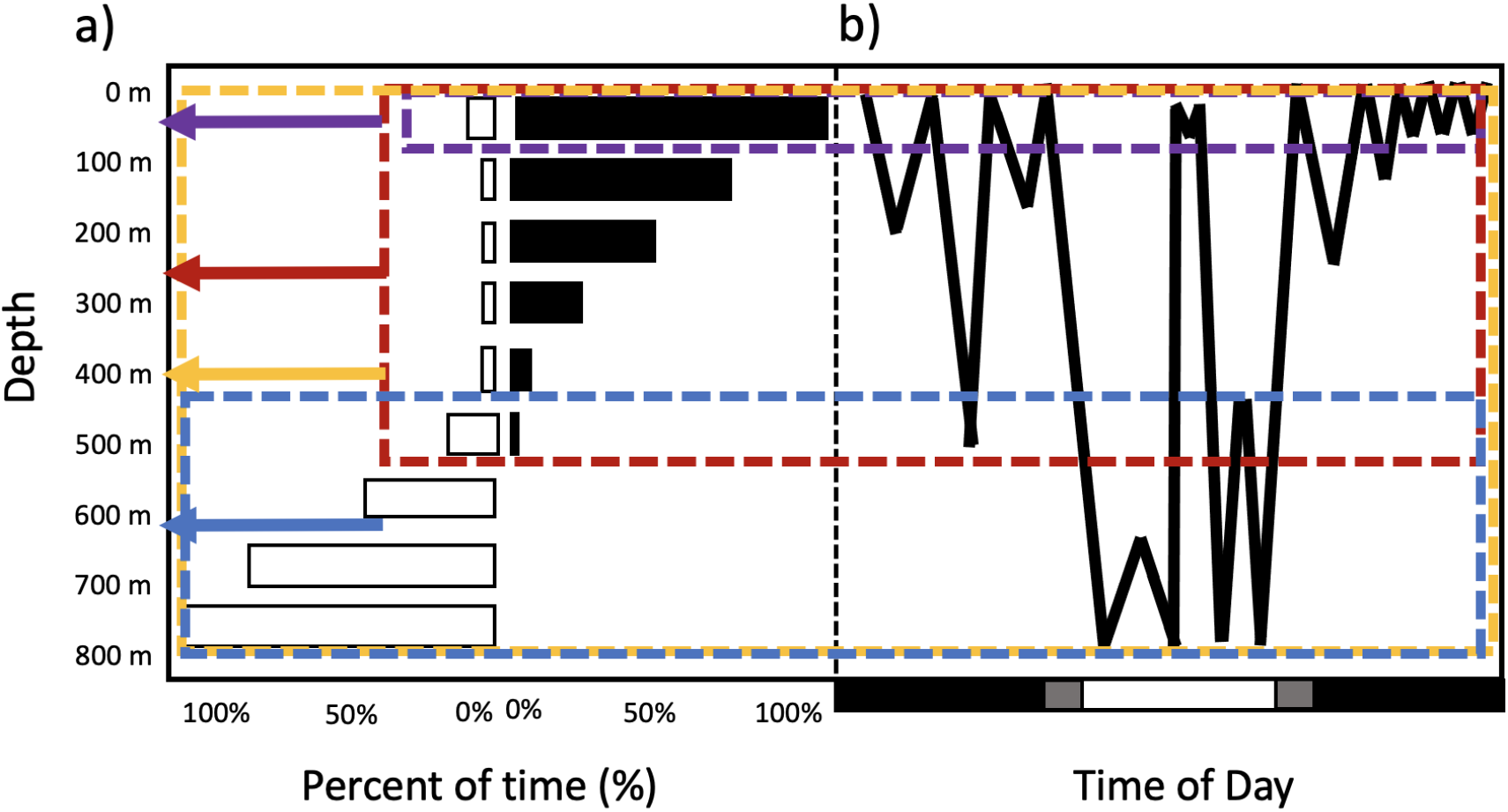
Illustration of typical tagging observations showing day and night vertical distribution of large marine predators, and extraction of preferred diving depth (D_pref_) and range (ΔD_pref_). (a) Histogram of the vertical distribution of a typical large marine predator. The white bars represent the proportion of time spent at a given depth during daytime, and the black bars during nighttime. (b) Vertical trajectory (i.e., depth as a function of time) for a typical large marine predator. The black, white, and gray bars at the bottom highlight nighttime, daytime and dawn/dusk periods respectively. In panels (a) and (b), the blue and purple boxes illustrate the extraction of the preferred diving depth during day and night respectively (here, D_pref_ ± ½ ΔD_pref_ = 600 m ± 200 m during the day, and 50 m ± 50 m at night). The yellow and red boxes represent the total range (ΔD = D_deep_-D_shallow_) of day and night diving depths (here, 0-800 m during the day, and 0-500 m at night).

We summarize these diverse data into two unambiguous quantities that could be estimated for both daytime and nighttime separately: (1) the preferred diving depth (D_pref_, blue and purple arrows in Fig. 1), representative of the approximate depth at which individuals spend most of the time; and (2) the preferred diving depth range (ΔD_pref_, blue and purple boxes Fig. 1), representative of the fraction of the water column where individuals are most commonly encountered. The preferred depth range can also be interpreted as a measure of the uncertainty for the preferred depth (allowing us to express it as D_pref_ ± ½ ΔD_pref_).

We used different approaches to estimate these two quantities empirically, based on the information presented in each study. When average diving depths were directly provided, these were taken as a measure of the mean preferred diving depth (D_pref_). If available, depth ranges or standard deviations around the mean were converted to preferred depth ranges (ΔD_pref_). When histograms and trajectories were shown as figures, we estimated D_pref_ and ΔD_pref_ by drawing a box around the depth range where predators were observed to spend most of their diving time (at least ∼⅔ of occurrence) and recorded the midpoint of the box (corresponding to D_pref_) and the distance between the upper and lower limits of the box (corresponding to ΔD_pref_).

Fig. 1a illustrates the extraction of the mean preferred diving depth and range based on histograms of the vertical distribution of a tagged predator. The blue dashed box shows the envelope of daytime depths, and includes most of the occurrences. In this scenario, the mean value was found by taking the midpoint of the range delimited by the blue box (i.e., D_pref_ = 600 m for the example of Fig. 1a), while the preferred range was defined as the distance between the upper and lower boundary (ΔD_pref_ = 400 m). The purple box shows the same approach for nighttime observations (i.e., D_pref_ ± ½ ΔD_pref_ = 50 m ± 50 m).

Fig. 1b illustrates the extraction of mean preferred diving depths and range based on vertical trajectories of the predator movement. Individual tracks show both day and night data on the same figure, separating day, night, and dawn/dusk periods. We disregarded dawn and dusk as transitional phases during which predators shift between nocturnal and diurnal behavior, and focused on observations during day and night only. The blue dashed box in Fig. 1b shows the range of preferred daytime depths and includes most of the occurrences, disregarding sporadic resurfacing when present. Once the depth range was identified, the midpoint of the box and upper/lower boundaries were used to determine the mean preferred diving depths (D_pref_) and preferred diving range (ΔD_pref_). A similar approach was applied to nighttime data (purple box in Fig. 1b). The behavior of LPP is sometimes affected by the tagging process, up to a day after attachment (Schaefer et al. 2009; Cartamil et al. 2010; Abascal et al. 2016). To account for this potential disturbance, we disregarded the early segments of vertical trajectories when the data reported allowed their identification.

For a handful of references, diving depths were reported under unique formats, e.g., as vertical density distributions as a function of time determined from multiple days (Queiroz et a. 2010). For these references, we applied a variation of the methods discussed above.

Together with information on preferred diving depths, we also extracted the total vertical range of a specific individual (see Fig. 1a, b, red and orange boxes). This is defined as the range between the deepest (D_deep_) and shallowest (D_shallow_) diving depths reported, and quantified by their difference (ΔD = D_shallow_ – D_deep_). While not shown in the main manuscript, these quantities inform our discussion of the results, and are presented in the Supplementary Information, Fig. S1 and S2.

### Ancillary information

Together with diving depth data, we extracted three additional types of information needed to co-locate diving depth observations with environmental variables, and to account for possible ontogenetic behavioral effects. These consist of the location and period of the observations, and the size of the tagged individual.

We estimated the location from the longitudes and latitudes of the horizontal tracks of tagged individuals (or groups of individuals) when they were provided. Fig. 2 illustrates the extraction of this spatial information. Similar to diving depths, we used horizontal tracks to determine spatial range boundaries. From these boundaries, we defined the midpoint coordinates of the area visited by the individual during the tagging period, x_Lon_ and x_Lat_, and the ranges around this midpoint, Δx_Lon_ and Δx_Lat_. When limited spatial information was provided, we used reported tagging or popup coordinates, or any regional information, to approximate representative area midpoints and ranges.

**Figure 2.**
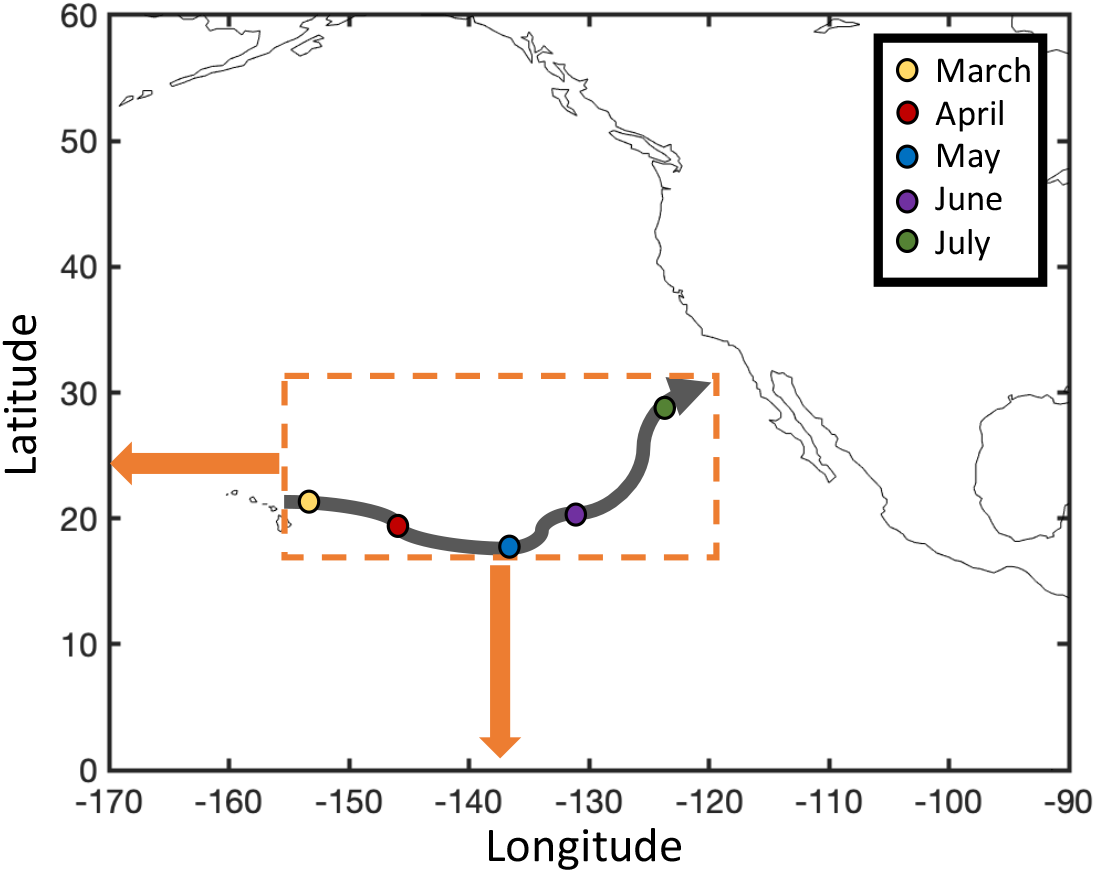
Illustration of typical observations of the horizontal range of a tagged large marine predator. The solid black line shows the track (longitude vs. latitude) of an individual predator, with the colored dots showing the beginning of each month. The box indicates the spatial range occupied by the individual during the period of tagging (here, 5 months), summarized by the average latitude and longitude and their ranges (here, x_Lat_ ± ½ Δx_Lat_ = 25 ± 7 N and x_Lon_ ± ½ Δx_Lon_ = 142 ± 18 W respectively).

To estimate the period based on depth versus time trajectories (see Fig.1b), we used the day, month and year at the beginning and end of tagging, which were provided in most studies. For histograms (see Fig.1a), the corresponding day, month and year at the beginning and end of the tagging period were generally provided, either directly, or in tables listing tagging and popup dates. When histograms were shown for multiple individuals, we used the tagging date of the earliest tagged individual and the latest popup date. When tagging occurred over several years but described the same seasons (or set of months), we recorded the starting and ending months of the period. When entire years were represented, we recorded the years and assumed that the data represented an average behavior over that period.

Finally, for almost every study, the size of the individual (or group of individuals) was provided, either as length or weight. We collected and converted these quantities to lengths in centimeters by applying standard conversion factors (see Supplementary Table S3 for conversions). We averaged representative sizes to the best of our understanding when diving depths applied to groups of LPP rather than single individuals, for example when data for juveniles vs. adults (Coelho et al. 2015), or males vs. females (Chapman et al. 2007) were reported. We normalized all lengths to fork length (L_FL_).

### Quality control and merged dataset

For each diving depth, we provided a quality ranking that reflected the confidence in the extraction process. When preferred diving depths were explicitly provided as numerical values, a rank “A” was assigned to the data points (30% of observations). When depths were inferred from figures but were clearly identifiable, a rank “B” was used (37%). Finally, when reasonable doubts about the accuracy of the estimates remained, a rank “C” was used (33%).

We replicated the data extraction independently three times. Two analysts (A. Nuno and J. Guiet) separately processed the full list of studies, while the third analyst (B. Baranek) randomly processed 20% of the studies for independent verification. Results from the three independent extractions show similar distribution of diving depths and ranges with only minor biases (Supplementary Information Fig. S3), suggesting robustness for our approach. Different extractions occasionally show shifts in diving depths (D_pref_), but always preserve relative depth variations. Sizes (L_FL_) are essentially identical across extractions since they are explicitly reported. Time and geographical locations show only minor spreads.

Quality rankings assigned by the two full data extractions were identical for “A” rank data points, and generally similar for “B” and “C” rank observations. Conservatively, we conducted the analysis of diving depths only on data classified as “A” or “B”, for which ranks from the two full extractions agreed. For the rest of the paper, we use the arithmetic mean of the two full extractions (analysts 1 and 2) for further analysis.

### Co-located environmental drivers

Environmental drivers of diving behavior generally reflect physiological constraints, for example water column temperature and oxygen (Patterson et al. 2008; Weng et al. 2009; Nasby-Lucas et al. 2009; Chiang et al. 2011), and behavioral constraints, such as the presence of prey (Sims et al. 2005; Vetter et al. 2008; Schaefer and Fuller 2010; Dewar et al. 2011; Braun et al. 2019). To parse key drivers of diving depth across a diverse range of species and regions, we co-located individual diving observations with climatological environmental information extracted from large-scale oceanographic databases and reanalysis products. For each diving observation, these environmental variables, listed in Table 1, were averaged over the spatial region of observations (x_Lon_ ± ½ Δx_Lon_ and x_Lat_ ± ½ Δx_Lat_), for the same climatological period.

**Table 1.** Selected drivers of diving behavior, classified into variables related to physiology, predator-prey interactions, and ontogeny. Environmental variables are co-located with diving observations from tagging studies (*Methods*). See Supplementary Fig. S4 for annual mean maps of the environmental variables.

| Category | Driver | Unit | Source | Resolution | Reference |
| --- | --- | --- | --- | --- | --- |
| Physiology | $\Delta O_2$ | $\mu\text{mol kg}^{-1}$ | Computed from O2 World Ocean Atlas 2018 | 1 degree & monthly | (Garcia et al. 2019) |
| | $\Delta T$ | $^{\circ}\text{C}$ | Computed from World Ocean Atlas 2018 | $\frac{1}{4}$ degree & monthly | (Locarnini et al. 2018) |
| | SST | $^{\circ}\text{C}$ | World Ocean Atlas 2018 | $\frac{1}{4}$ degree & monthly | (Locarnini et al. 2018) |
| Predator-prey | $Z_{eu}$ | m | Mean of 3 reconstructions (CbPM/VGPM/Eppley-VGPM) | $\frac{1}{6}$ degree & monthly | (Behrenfeld and Falkowski 1997; Carr et al. 2006; Marra et al. 2007) |
| | $Z_{meso}$ | m | Reconstruction | 2 degrees & monthly | (Bianchi et al. 2013; Bianchi and Mislán 2016) |
| | Chl | $\text{mg m}^{-3}$ | Climatology computed from SeaWiFS observation (1998-2004) | 1 degree & monthly averaged | (SeaWiFS) |
| | EKE | $(\text{m/s})^2$ | Computed from AVISO velocity anomalies (2000-2018) | $\frac{1}{4}$ degree & monthly averaged | (AVISO) |
| Ontogeny | $L_{FL}$ | cm | Converted to FL from references | - | - |

For poikilotherm species, water temperature modulates physiological rates. To evaluate temperature effects, we use climatological sea surface temperature (SST), and the climatological vertical temperature gradient (ΔT) between the surface and 250 m depth, using the World Ocean Atlas 2018 (WOA 2018, Locarnini et al., 2018) (Table 1; see maps in Supplementary Fig. S4).

For some species, behavioral and physiological adaptations mitigate the effects of temperature variations during deep dives. Among these, one of the most effective is the *rete mirabile*, a circulatory apparatus that maintains higher temperatures for selected organs in the predator’s body. These include brain, eyes, muscles, kidneys, and stomachs, enhancing prey detection, swimming, and digestion respectively (Burne 1924; Block 1986; Brill 1994; Weng and Block 2004; Fritsches et al. 2005; Stoehr et al. 2020). We approximate the degree of thermoregulation by quantifying the number of physiological adaptations for each species (Supplementary Table S3). Temperature sensitivities may also vary with life history and individual size (Hino et al. 2019), here encapsulated by L_FL_.

To characterize the role of oxygen on deep dives, we extracted the oxygen gradient (ΔO_2_) between the surface and 250 m depth, as a proxy for the depth of low-oxygen layers (Table 1), based on climatologies from the WOA 2018. To assess for the role of prey occurrence, we use reconstructions of the daytime depth of mesopelagic sound scattering layers (Z_meso_) from a global analysis of acoustic observations (Bianchi et al., 2013; Bianchi & Mislan, 2016, see Table 1). To consider the effect of irradiance in the water column, we use the depth of the euphotic zone (Z_eu_) as a proxy of light penetration (Table 1). We also compare diving depths with quantities known to affect lower trophic levels (Arostegui et al. 2022), including chlorophyll-a concentrations (Chl) from remote-sensing products, and eddy kinetic energy (EKE) from AVISO surface velocities (Table 1).

Some of the predators considered here are opportunistic feeders, while others are more selective. For instance, swordfish and bigeye thresher sharks feed on a variety of epipelagic and mesopelagic species (Preti et al. 2008), while bigeye tuna are more closely associated with vertically migrating mesopelagic fish (Ménard et al. 2000). To account for the effects of prey type and diversity, we estimated the flexibility of the diet of LPP by quantifying the number of functional groups targeted by each species (Supplementary Table S3), based on information from FishBase (Froese and Pauly 2022).

## Results

### Global distribution and patterns of diving depth

We extracted a total of 671 independent observations of preferred diving depth from 24 different species. From this point forward we focus on 21 species for which at least 3 observations were obtained. Of these, 85% include explicit day and/or night information, with 78% ranked as A or B in quality. Observations span the global ocean (Fig. 3), ranging from coastal to open waters, and are more abundant in the North Pacific Ocean, North Atlantic Ocean, and around Australia. Most are from subtropical and temperate latitudes, with patterns reflecting the typical range of each species. While billfish observations come mostly from subtropical regions, tuna and sharks are commonly sampled at higher latitudes (Fig. 3a).

**Figure 3.**
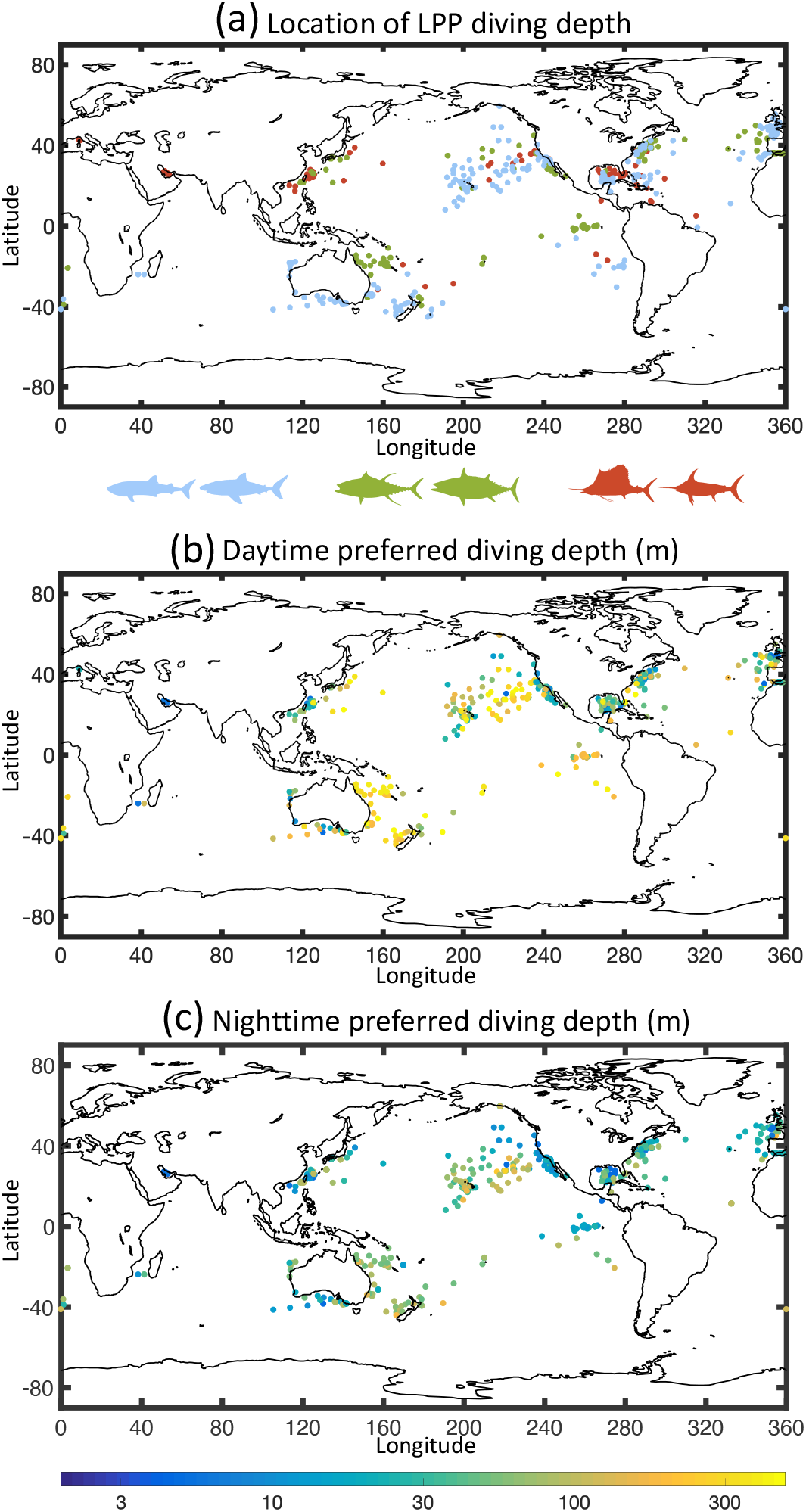
Spatial distribution and diving depths of tagged large marine predators. (a) Average position of each diving depth observation from a meta-analysis of published studies. Colors indicate three broad taxonomic groups (blue=sharks; green=tuna; red=billfish), for shallow and deep diving species. (b) Preferred daytime diving depths (m) for each data point in (a). (c) Preferred nighttime diving depths (m).

Coherent, large-scale patterns in diving depths (D_pref_) are apparent (Fig. 3b,c), with predators generally diving deeper offshore and during the daytime, and shallower closer to the coast and in productive waters, and during the nighttime. A large cluster of predators observed in the open ocean between the West Coast of North America and Hawaii, and around Galapagos, show intermediate to deep daytime diving depths (>300 m). In coastal waters along Eastern Australia, we observe a cluster of predators with relatively deep diving behavior (300-400 m). In coastal waters along North America, Southwest Australia, Western Europe and Southeast Asia, predators tend to dive shallower (<100 m). At the large scale, the type of predator appears to play a secondary role for the diving depth, suggesting common environmental constraints across species (compare Fig. 3a and b-c). Nonetheless, patchiness in diving depths within each region also indicate significant species-dependent variability.

### Preferred diving depths

Across the 21 species sampled, most pelagic predators dive deeper during the day than during the night (Fig. 4), except for oceanic whitetip sharks, whale sharks, and sailfish, for which nighttime depths are approximately 10 m deeper than daytime depths on average. Two main categories of diving behavior emerge: (1) deep diving species with much deeper daytime preferred depths that reach well within the mesopelagic zone (deeper than 150 m), and (2) shallow diving species with slightly deeper daytime preferred depths that generally remain in the epipelagic zone (shallower than 150 m). Bigeye thresher sharks show the largest day to night difference, up to 280 m. For shallow divers, diurnal variations are less pronounced, and day/night differences are less than 50 m, with a maximum for white marlin. Each taxonomic group includes species that belong to the deep or shallow diving categories. All billfish except swordfish belong to shallow divers.

**Figure 4.**
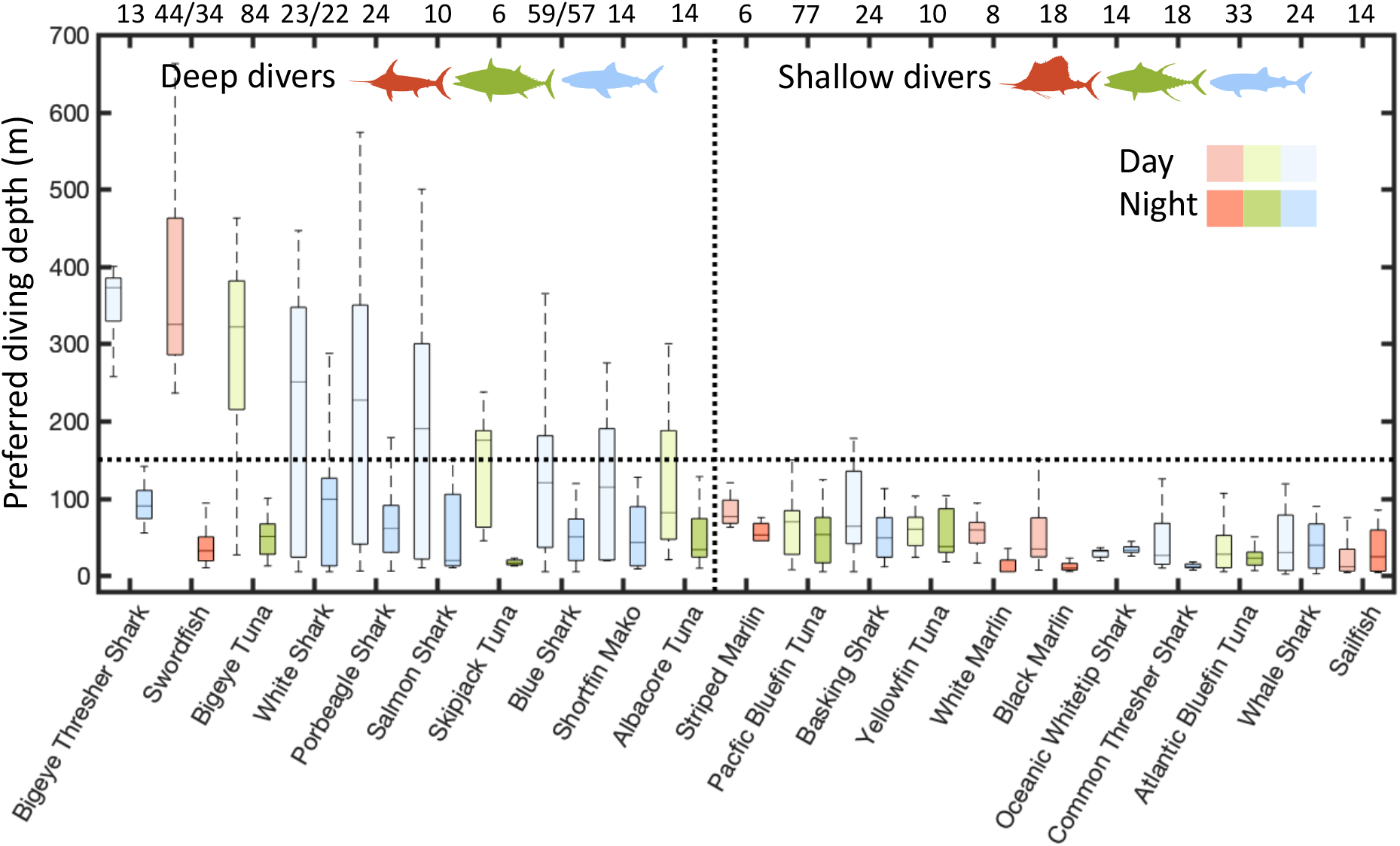
Daytime and nighttime preferred diving depth (D_pref_) by species and taxonomic group. Results are shown for all data points, omitting three species with less than 3 data points each (blue marlin, spearfish, and southern bluefin tuna). In each box plot, the central mark indicates the median, the top and bottom edges the 75^th^ and 25^th^ percentiles, and the whiskers the full range. Species are ranked from deepest to shallowest based on daytime preferred depths. The number of observations for each species is shown on top (day/night). Taxonomic groups are shown by different colors (blue=sharks; green=tuna; red=billfish) for daytime (light shades) and nighttime (dark shades). See Supplementary Fig. S1 for a similar figure showing the maximum recorded diving depths (D_deep_).

Variability in preferred diving depths does not perfectly match the variability in the maximum diving depth (D_deep_) across species (Supplementary Fig. S1). Some shallow diving predators complete deep sporadic dives (>500 m), especially bluefin tuna, whale sharks, and basking sharks. However, most shallow diving species remain in the ocean epipelagic layers.

Preferred depths ranges vary systematically by taxonomic group and between deep and shallow divers, with variable day/night overlaps (Fig. 5). Among deep diving LPP, nighttime diving depths are similar across groups (see Fig. 5a-c), with somewhat deeper depths for sharks, and shallower depths for swordfish. In contrast, daytime depths are more variable across deep diving groups. During the day, the overlap between shallow and deep dives is more pronounced for sharks, which, while occasionally venturing in relatively deep waters (>400 m depth) also spend a significant amount of time near the surface (Fig. 5a). For tuna, the daytime distribution is deeper on average, with a peak in the 200-400 m depth range (Fig. 5b). A significant occurrence in shallower waters also suggests frequent excursions to the surface. For swordfish, daytime and nighttime habitats are nearly completely disconnected, with typical daytime depths in the 300-600 m range, and limited surface occurrences. These systematic differences suggest distinct deep diving behavior across groups. Within each taxonomic group, day/night diving depths vary by species, with the deepest divers inhabiting increasingly disconnected layers between day and night (Supplementary Fig. S5).

**Figure 5.**
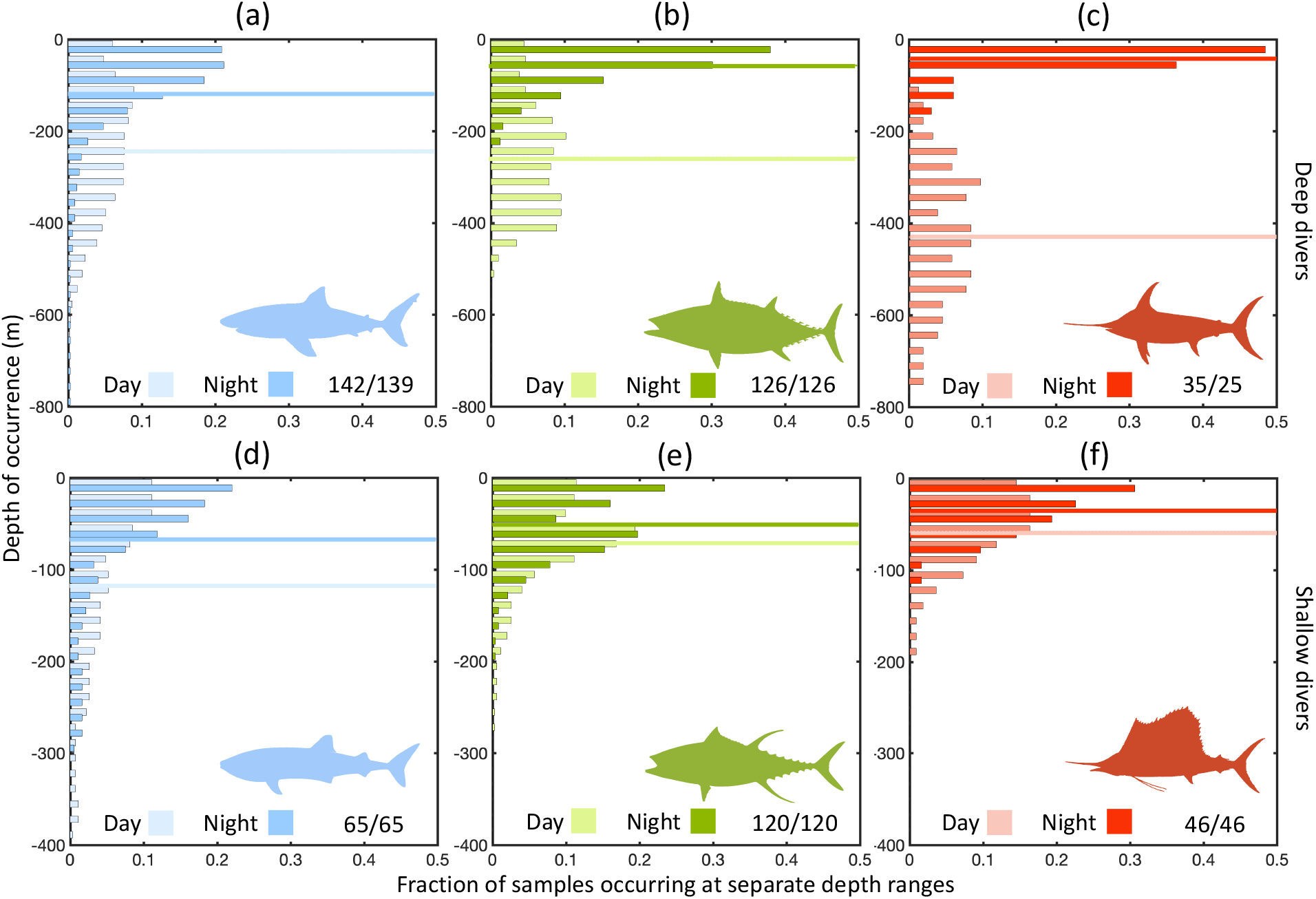
Distribution of day and night preferred vertical depths (D_pref_) for large marine predators based on tagging data. Observations are separated by taxonomic group (blue=sharks; green=tuna; red=billfish). The top row (a-c) shows deep diving species, and the bottom row (d-f) shallow diving species. In each panel, the horizontal line shows the mean preferred depth, and the numbers the total number of samples (day/night). See Supplementary Fig. 5 and 6 for similar observations grouped by species. Note the different vertical ranges for deep and shallow diving species.

For shallow diving LPP, nighttime/daytime preferred depths are similar across taxonomic groups. All occur between the surface and 150 m during day and night (Fig. 5d-f), with only a slight deepening by up to 50 m during the day. A deeper, infrequent daytime peak at around 350 m for sharks (Fig. 5d) can be attributed to a single species, basking sharks (see Supplementary Fig. S6), which occasionally perform diel vertical migrations into the mesopelagic ocean (Dewar et al. 2018). Considering these sporadic deep dives, sharks dive slightly deeper on average, followed by tuna and billfish, both during daytime and nighttime.

### Correlations with environmental variables

To identify potential drivers of diving behavior, we analyze the correlations between day and night preferred depths and co-located environmental variables (Fig. 6). We also assess the significance of the combined selected drivers by comparing the adjusted coefficient of determination of a multilinear regression between all drivers and the observed diving depths (last columns in Fig. 6a,b; see Supplementary Table S4 for the parameters of the multilinear regressions).

**Figure 6.**
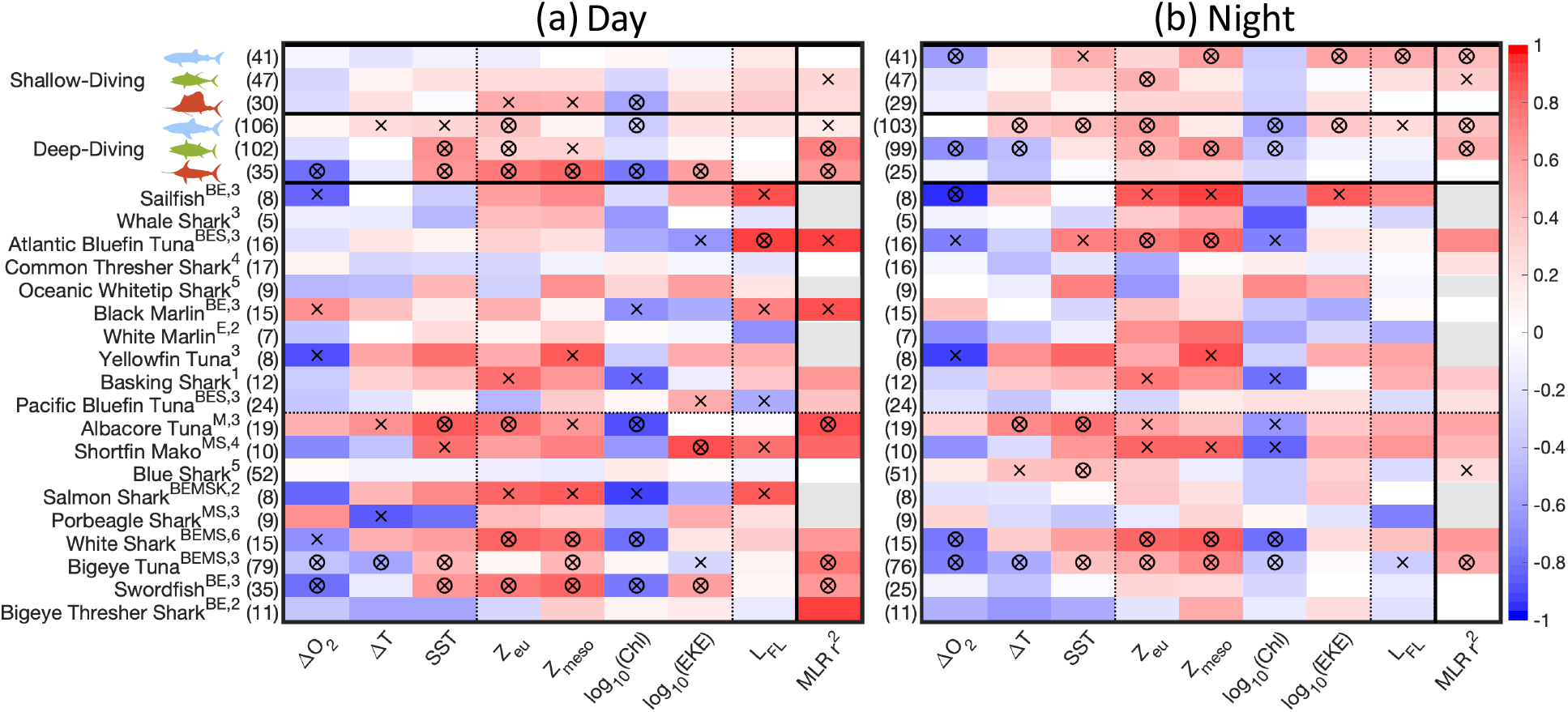
Correlation between diving depths of large marine predators (D_pref_) and selected environmental drivers for (a) daytime and (b) nighttime. Colors show the Pearson correlation coefficient r between individual drivers and D_pref_ for each category (red=positive correlation; blue=negative correlation). In each panel, the leftmost column shows the adjusted r^2^ of the multilinear regression between all drivers and D_pref_. Gray boxes show cases with too few observations to conduct a multiple linear regression. Crosses and circles indicate correlations at the 1% and 0.1% significance levels respectively. The top rows show correlations for aggregated taxonomic groups (blue=sharks; green=tuna; red=billfish), separating shallow from deep diving species. The bottom rows show correlations for individual species, ranked from shallowest to deepest. For each species, the letters in superscript indicate the presence of thermoregulation apparatuses (B=brain, E=eyes, M=muscles; S=stomach; K=kidney; see also Supplementary Table S3). The numbers in superscript indicate the degree of feeding generalism (i.e., the number of different prey functional groups, see Supplementary Table S3). Correlations are computed for data ranked A and B in quality, removing outliers and disregarding species with less than 3 samples each (blue marlin, spearfish, and southern bluefin tuna). See Supplementary Fig. S2 for a similar figure based on maximum diving depths (D_deep_).

Across all species, euphotic zone depth, sound scattering layer depth, and surface chlorophyll show consistent significant correlations with day and night diving depths (Fig. 6a,b). LPP generally dive shallower in high-chlorophyll, productive regions, and deeper in regions with low chlorophyll, increased light penetration, and deeper sound scattering layers. Surface temperature also shows a generally consistent correlation across species, with LPP diving deeper under warmer SST. Similarly, stronger vertical oxygen gradients correlate more frequently with shallower dives. Other drivers show less consistent and less robust correlations. For instance, vertical temperature gradients and EKE show a mixed influence across groups and species. Often, larger predators tend to dive deeper than smaller predators, although predator size alone is overall a poor predictor of diving depth.

For shallow diving species, most environmental drivers show a weak influence on diving behavior during the day, except for drivers related to predator-prey interactions (Z_eu_, Z_meso_, Chl) for billfish. A multilinear regression considering all drivers has weak explanatory power (r^2^ up to 0.27 for tuna). At night, correlations are stronger, especially for sharks, with higher coefficients of determination (r^2^>0.35 for shark and tuna).

In contrast, for deep diving species, preferred diving depths are more strongly correlated with environmental drivers, especially during the day (r^2^ up to 0.73 for tuna), but also at night for shark and tuna (r^2^>0.4). Drivers related to predator-prey interactions (Z_eu_, Z_meso_, Chl, EKE) are especially significant during the day, but also at night for shark and tuna. Drivers linked to physiology (ΔO_2_, ΔT and SST) are also significant, in particular for tuna and swordfish during the day, and for shark and tuna at night, but with somewhat more mixed influences.

Correlations by diving type or taxonomic group show emergent relationships with selected drivers. Shallow diving sharks and tuna are more strongly influenced by environmental drivers at night relative to the day. In contrast, diving depths of shallow and deep diving billfish show significant correlations with environmental variables during the day, but not at night. Deep diving sharks and tuna generally show stronger correlations for both day and night observations. However, these emergent relationships are remarkably variable on a species-by-species case, with stronger correlations for deeper divers, likely reflecting physiological adaptations to vertical temperature gradients (e.g., the *rete mirabilis*) and prey preference. Consistently, the deepest diving predators can maintain uniform internal temperatures across multiple organs, while shallow diving predators, especially sharks, generally lack thermoregulation apparatuses (Fig. 6).

Correlations between preferred diving depths and environmental drivers are similar when using the maximum diving depth (D_deep_) rather than the preferred depth (see Supplementary Fig. S2), suggesting common patterns and drivers for the two quantities.

## Discussion

Despite some overarching commonalities, systematic differences in diving behavior between shallow and deep divers, and, within these categories, between shark, tuna, and billfish, suggest that selected drivers affect diving behavior differently across species.

### Deep divers

For deep diving species, access to the mesopelagic prey is a main driver of diving behavior. This is supported by large day-night differences in vertical habitat that match the day-night excursion of vertically migrating mesopelagic nekton (Fig. 4). It is also supported by significant correlations with the depth of sound scattering layers (Z_meso_ in Fig. 6). This association has been documented for deep diving tuna, swordfish, and sharks (Dagorn et al. 2000; Schaefer and Fuller 2002; Abascal et al. 2010; Campana et al. 2011; Williams et al. 2015; Francis et al. 2015b). Weaker correlations for the preferred depth of deep diving sharks and Z_meso_ might reflect large intra-species variations in vertical habitat use, for example between traveling and foraging phases (Nasby-Lucas et al. 2009), shifts in behavior from coastal to offshore regions (Bonfil et al. 2010; Queiroz et al. 2010), or diets less dependent on mesopelagic fish. This is supported by the broader spread of diving depths for shark species (Fig. 5a,d)

Significant correlations with proxies of light penetration (Z_eu_) and prey abundance (Chl) support the importance of visual predation for deep divers, but the underlying mechanisms remain unclear. Light penetration is correlated to both surface chlorophyll concentration and the daytime depth of vertically migrating mesopelagic nekton (Bianchi et al. 2013; Bianchi and Mislan 2016; Klevjer et al. 2016), making it difficult to separate the specific influence of these variables on predators or prey.

Correlations with variables that relate to metabolism (ΔO_2_, ΔT, SST) support a role for physiological constraints for some deep diving species. For sharks, weaker correlations with temperature suggest distinct vertical behaviors relative to other groups, with residence in the warm surface punctuated by sporadic ventures in deeper layers. This behavior contrasts to the more consistent, long-lasting diel vertical migrations to mesopelagic depths observed for deep diving tuna and swordfish (Evans et al. 2008; Abecassis et al. 2012). Oxygen vertical gradients are known to affect the duration of deep dives and the frequency of resurfacing (Dagorn et al. 2000; Hino et al. 2019). The mixed influence of vertical oxygen gradients shown by our analysis suggests that swordfish and tuna may be more sensitive to this variable than other species (Fig. 4).

Some species can also exploit patchiness in the physical environment. For example, blue sharks take advantage of warm anticyclonic eddies near the Gulf Stream to reach deeper layers (Braun et al. 2019), and eddies have been shown to affect the catch of many LPP in the subtropical North Pacific (Arostegui et al. 2022). Here, we find only a minor effect of EKE, an imperfect proxy of environmental heterogeneity.

Environmental drivers, in particular those that affect an organism’s physiology, could influence preferred diving depths indirectly, by affecting the distribution of prey, or directly, by impacting predator performance and feeding success. Disentangling these influences is challenging, especially with the type of aggregated data used in this study. However, some remarkable features should be noted. In general, deeper diving species, independently of taxonomy, show stronger sensitivities to drivers related to physiology (Fig. 6), and a stronger disconnection between daytime and nighttime vertical distributions (Fig. 4). A larger energetic cost for deeper dives might require longer foraging times at depth, and hence tighter physiological adaptations that may make these species more susceptible to the characteristics of the environment. For swordfish, we observe a complete loss of significant correlations with all drivers at night, while most drivers are highly significant during the day, when these species occupy the deepest layers. This suggests that swordfish may be released by physiological or performance constraints when they occupy the epipelagic layers at night. This contrasts with deep diving tuna and shark, for which correlations are somewhat stronger in shallower layers at night, relative to the day.

### Shallow divers

Shallow diving LPP show significant variations in preferred depths between day and night, but with much smaller amplitudes than deep diving species (Fig. 4). This indicates a preference for warmer, well lit, and well oxygenated surface waters. For these species, day to night differences are likely related to the occurrence of prey, or to environmental characteristics that facilitate prey capture, e.g., by making it more visible (Itoh et al. 2003; Pohlot and Ehrhardt 2018).

Temperature and oxygen have been shown to constrain the vertical habitat of some shallow diving species (Stramma et al. 2012). For shallow diving shark and tuna, the absence of significant correlations with environmental drivers during the day, and stronger correlations at night, suggest that when mesopelagic prey migrate to deep layers, factors not considered here may become important. These could include diurnal and finer-scale variation of illumination (Patterson et al. 2008; Omand et al. 2021), ocean currents (Kitagawa et al. 2004; Williams et al. 2017), occurrence of shallow prey (Horodysky et al. 2007), or ontogeny (Kitagawa et al. 2007; Cartamil et al. 2016). For billfish species, remarkable correlations with proxies of light availability during the day suggest an important role for visual predation.

We also note that, while our analysis focuses on the most common diving depths, some species classified as shallow divers, e.g., Atlantic bluefin tuna, whale sharks and basking sharks, may occasionally dive to mesopelagic depths, likely to forage on deep-dwelling prey (Brunnschweiler et al. 2009; Dewar et al. 2018).

### Other drivers of diving behavior

Our analysis is limited by the type of proximate environmental variables that could be co-located with tagging studies. However, additional key drivers are likely to play a significant role. For instance, some deep dives could help navigation of migrating LPP (Willis et al. 2009). These deep dives occur in the twilight hours and would only marginally influence the preferred day and night depth determined in our study. Similar correlations between environmental variables and maximum and preferred diving depths (Supplementary Fig. S2) suggest that both quantities reflect similar drivers.

Other behaviors not considered in this study influence intra-species variations in diving depths, including social interactions during mating (Jorgensen et al. 2012) and parasite removal (Braun et al. 2022). Different hunting strategies would also affect the shape of vertical migrations (Weng et al. 2007; Abascal et al. 2011). Finally, diving behavior is altered by human practices such as the use of fish aggregating devices that disrupt vertical migration patterns (Schaefer and Fuller 2002, 2010; Musyl et al. 2003), the release of bait in the water, or “chumming” (Hueter et al. 2018), and the tagging process itself (Sippel et al. 2011; Abascal et al. 2016).

Fine-scale variability not considered in this study could also be important. For example, the presence of fronts and eddies can alter the physical conditions in the water column (Lévy et al. 2018), as well as predator and prey behavior. While we tested the importance of EKE as a proxy of fine-scale variability (Fig. 6), its effect remains mixed. For species like blue shark, mesoscale eddies facilitate deep dives (Braun et al. 2019). Fronts and eddies can also drive prey aggregation at the surface, altering predator behavior, as suggested for basking sharks (Sims et al. 2005; Shepard et al. 2006). A widespread influence of mesoscale eddies on a variety of LPP was also suggested by longline fisheries data in the North Pacific Ocean (Arostegui et al. 2022), likely reflecting effects on the distribution of prey. Similarly, variations in surface light may modulate diving behavior (Matsumoto et al. 2013). These include rapid changes caused by clouds during the day (Omand et al. 2021), or different lunar phases at night (Abascal et al. 2010; Dewar et al. 2011; Tanaka and Yamaguti 2017).

### Caveats

To minimize uncertainties in the extraction of diving depths, we followed an established meta-analysis approach (Haddaway et al. 2020), which we replicated independently twice with all the available data, and a third time on a subset of the data for verification. To address the sources of uncertainty related to the way diving data is reported in different studies, we ranked the quality of our extractions, using only reliable data points (ranked as A and B) for the final analysis. Importantly, our results are robust to the inclusion of data of poorer quality (ranked as C) (see Supplementary Fig. S7 and S8). However, when only A ranked data are considered, many correlations lose significance, likely because of the reduction in the number of data points.

Our analysis also relies on data aggregation on multiple spatial and temporal scales (days to years). Since seasonal and spatial variations affect diving behavior, binning data on coarse spatial and temporal scales likely reduces our ability to detect significant patterns. To test the effect of data aggregation, we repeated the correlation analysis (Fig. 6) with observations representative of small (<10^5^ km^2^) or large (>10^5^ km^2^) oceanic regions, and of seasonal (<4 months) or annual (>4 months) periods (Supplementary Fig. S7 and 8). This analysis indicates that the emergent correlations discussed in this paper are conserved at different levels of aggregation.

## Conclusions

We conducted a meta-analysis of diving behavior for 24 species of LPP from tagging data, based on the extraction of 671 individual diving depth observations from 87 studies. At regional scales, common behaviors emerge for all species, with deeper dives in the open ocean, and shallower dives in productive coastal regions. By co-locating individual diving depths with environmental drivers, we also uncover group- and species-specific influences on diving behavior.

The analysis shows that LPP can be divided into two main categories, shallow and deep divers, irrespectively of their broad taxonomic characterization. Deep diving species are influenced by the depth of mesopelagic prey, and a combination of light penetration and the productivity of surface waters. Accordingly, we observe deeper dives in clearer waters, where mesopelagic sound scattering layers are found at greater depths in the water column. Deep diving species are also influenced by variables that impose physiological constraints to predator activity, with deeper dives in more oxygenated, warmer waters. However, the links between metabolic drivers and diving depths of LPP remain unclear, because they likely influence the diving behavior of both predators and prey simultaneously.

In general, diving depths are more predictable for deep-diving species during the day, suggesting tighter physiological adaptations or prey specificity. For shallow diving species, weaker correlations with environmental drivers suggest more variable diving behavior, especially during the day, and a stronger influence of fine-scale heterogeneity in the surface habitat, e.g., caused by changes in light or mesoscale currents, which are hard to capture with climatological data.

The overarching patterns highlighted by our analysis open the door for a mechanistic, quantitative characterization of physiological and ecological constraints on vertical habitat use for a variety of LPP species. They also highlight the challenge of disentangling multiple overlapping influences on predator and prey behavior. Future work based on direct analysis of vertical tracks for tagged species, rather than on meta-analysis, will likely reduce the uncertainty of our results, establishing tighter constraints on the relationships between the environment and diving depths. In turn, a more quantitative understanding of the patterns and drivers of LPP diving behavior will facilitate detection of changes in the vast oceanic ecosystem, support management and protection of valuable species, and improve marine ecosystem models for climate change and biogeochemical studies.

## Acknowledgments

This material is based upon work supported by the California Ocean Protection Council under Grant No. C0100400, and the National Aeronautics and Space Administration (NASA) under Grant No. 80NSSC21K0420 issued through the Ocean Biology and Biogeochemistry (OBB) program. DB acknowledges support from the Alfred P. Sloan Foundation. Computational resources were provided by the Expanse system at the San Diego Supercomputer Center through allocation TG-OCE170017 from the Extreme Science and Engineering Discovery Environment (XSEDE), which was supported by National Science Foundation grant 1548562.

