## Supplementary Information for "Patterns and drivers of the diving behavior of large pelagic predators"

**Table S1: List of keywords searched through the Web of Science and Google Scholar. We sequentially looked for publications in the three categories "migration", "migration + classes" and "migration + specific species".**

| *Categories* | *Keywords* |
|---|---|
| Migration | Vertical Migration; Diel Vertical Migration; Diving Depth; Vertical Behavior; Vertical Movements. |
| Classes | Shark; Billfish; Tuna; Pelagic Predators; Pelagic Sharks; Pelagic Billfish; Pelagic Tuna. |
| Specific species | Bigeye Tuna; White Shark; Bigeye Thresher Shark; Porbeagle Shark; Swordfish; Salmon Shark; Skipjack Tuna; Blue Shark; Shortfin Mako; Albacore Tuna; Striped Marlin; Basking Shark; Yellowfin Tuna; White Marlin; Black Marlin; Oceanic Whitetip Shark; Common Thresher Shark; Atlantic Bluefin Tuna; Whale Shark; Sailfish. |

**Table S2: List of papers considered for data extraction for 24 pelagic species of shark, tuna and billfish. The type of information provided is specified (H, histogram; P, profile; N, numeric value; O, other).**

| *Species* | *References* | *Type of data* |
|---|---|---|
| Albacore Tuna | (Childers et al. 2011)<br>(Cosgrove et al. 2014)<br>(Williams et al. 2015) | H/P/N<br>H/P/N<br>H/P/N |
| Atlantic Bluefin Tuna | (Wilson et al. 2005)<br>(Walli et al. 2009)<br>(Galuardi and Lutcavage 2012)<br>(Abascal et al. 2016) | H/P<br>P<br>H/N<br>P |
| Basking Shark | (Skomal et al. 2004)<br>(Sims et al. 2005)<br>(Shepard et al. 2006)<br>(Curtis et al. 2014)<br>(Braun et al. 2018)<br>(Dewar et al. 2018)<br>(Doherty et al. 2019) | H<br>P<br>P<br>H<br>N<br>H/P<br>H |
| Bigeye Thresher Shark | (Nakano et al. 2003)<br>(Weng and Block 2004)<br>(Stevens et al. 2010)<br>(Musyl et al. 2011)<br>(Coelho et al. 2015) | P<br>H<br>H<br>N<br>N |
| Bigeye Tuna | (Dagorn et al. 2000)<br>(Schaefer and Fuller 2002)<br>(Musyl et al. 2003)<br>(Arrizabalaga et al. 2008)<br>(Evans et al. 2008)<br>(Schaefer et al. 2009)<br>(Schaefer and Fuller 2010)<br>(Matsumoto et al. 2013)<br>(Hino et al. 2019) | P<br>H<br>N<br>P<br>H/N<br>H/P<br>H/P<br>H/P<br>P |
| Black Marlin | (Chiang et al. 2015)<br>(Williams et al. 2017) | H/N<br>P |
| Blue Marlin | (Goodyear et al. 2008)<br>(Kraus and Rooker 2007) | H<br>P |
| Blue Shark | (Carey et al. 1990)<br>(Klimley et al. 2002)<br>(Queiroz et al. 2010)<br>(Stevens et al. 2010)<br>(Campana et al. 2011)<br>(Musyl et al. 2011)<br>(Queiroz et al. 2012)<br>(Heard et al. 2018) | P<br>H<br>O<br>H<br>H<br>N<br>H<br>H |
| Common Thresher Shark | (Cartamil et al. 2010)<br>(Stevens et al. 2010)<br>(Cartamil et al. 2011)<br>(Cartamil et al. 2016) | H/P<br>H<br>N/O<br>H/P |
| Oceanic Whitetip Shark | (Musyl et al. 2011)<br>(Howey-Jordan et al. 2013) | N<br>H/N |
| Pacific Bluefin Tuna | (Marcinek et al. 2001) | H/P |

| | | |
|---|---|---|
| | (Itoh et al. 2003)<br>(Kitagawa et al. 2004)<br>(Kitagawa et al. 2007)<br>(Furukawa et al. 2017) | H/P<br>N<br>N<br>N |
| Porbeagle Shark | (Pade et al. 2009)<br>(Francis et al. 2015) | H<br>H/P |
| Sailfish | (Hoolihan and Luo 2007)<br>(Chiang et al. 2011)<br>(Hoolihan et al. 2011)<br>(Kerstetter et al. 2011)<br>(Pohlot and Ehrhardt 2018) | H<br>H<br>P<br>H/P<br>P |
| Salmon Shark | (Carlisle et al. 2011)<br>(Coffey et al. 2017) | P<br>H |
| Shortfin Mako | (Klimley et al. 2002)<br>(Loefer et al. 2005)<br>(Vetter et al. 2008)<br>(Stevens et al. 2010)<br>(Abascal et al. 2011)<br>(Musyl et al. 2011) | H<br>H/P<br>P<br>H<br>H/N<br>N |
| Skipjack Tuna | (Schaefer and Fuller 2007)<br>(Schaefer et al. 2009) | H/P<br>P |
| Southern Bluefin Tuna | (Patterson et al. 2008) | P |
| Spearfish | (Arostegui et al. 2019) | H/P |
| Striped Marlin | (Brill et al. 1993)<br>(Sippel et al. 2011) | P<br>H |
| Swordfish | (Takahashi et al. 2003)<br>(Abascal et al. 2010)<br>(Sepulveda et al. 2010)<br>(Dewar et al. 2011)<br>(Abecassis et al. 2012)<br>(Evans et al. 2014)<br>(Tanaka and Yamaguti 2017)<br>(Sepulveda et al. 2018) | H/P<br>H/N<br>H/P/N<br>H/P<br>P<br>H/P<br>H<br>H/N |
| Whale Shark | (Graham et al. 2006)<br>(Wilson et al. 2006)<br>(Brunnschweiler et al. 2009)<br>(Tyminski et al. 2015) | P<br>H/P<br>H/N<br>H/N |
| White Marlin | (Horodysky et al. 2007)<br>(Hoolihan et al. 2015)<br>(Vaudo et al. 2018) | H/P/N<br>H<br>H |
| White Shark | (Klimley et al. 2002)<br>(Weng et al. 2007a)<br>(Weng et al. 2007b)<br>(Nasby-Lucas et al. 2009)<br>(Bonfil et al. 2010)<br>(Jorgensen et al. 2012) | H<br>H/O<br>P/N/O<br>H/P<br>H<br>P |
| Yellowfin Tuna | (Block et al. 1997)<br>(Schaefer et al. 2009)<br>(Weng et al. 2009)<br>(Weng et al. 2017) | H<br>P<br>H<br>H/N/O |

**Table S3: Species specific information for the 21 main species considered in the study. Conversion factors for size normalization to fork length (FL) and for weight (KG) normalized to total length (TL) and feeding habits are based on FishBase (Froese and Pauly 2022). Organs with thermoregulation apparatus identified from literature.**

| *Species* | *Conversion size vs. weight* | *Feeding habits* | *Thermoregulation adaptations* |
|---|---|---|---|
| Albacore Tuna | FL = TL/1.083<br>TL= $(KG*1000/0.01862)^{(1/2.99)}$ | fish, invertebrates, crustaceans | muscle |
| Atlantic Bluefin Tuna | FL = TL / 1.075<br>TL= $(KG*1000/0.01259)^{(1/3.01)}$ | fish, invertebrates, crustaceans | brain, eyes, stomach |
| Basking Shark | FL = 0 + 0.909 x TL<br>TL= $(KG*1000/0.00389)^{(1/3.12)}$ | plankton | |
| Bigeye thresher shark | FL = 4.83 + 0.580 TL<br>TL= $(KG*1000/0.01096)^{(1/2.91)}$ | fish, invertebrates | brain, eyes |
| Bigeye Tuna | FL = 0 + 0.913 x TL<br>TL= $(KG*1000/0.01349)^{(1/3.02)}$ | fish, invertebrates, crustaceans | brain, eyes, muscle, stomach |
| Black Marlin | FL = TL / 1.127<br>TL= $(KG*1000/ 0.00447)^{(1/3.13)}$ | fish, invertebrates, crustaceans | brain, eyes |
| Blue Shark | FL = 1.391 + 0.831 x TL<br>TL= $(KG*1000/0.00447)^{(1/3.11)}$ | fish, invertebrates, crustaceans, sharks, birds | |
| Common Thresher Shark | FL = 7.0262 + 0.547 x TL<br>TL= $(KG*1000/0.00832)^{(1/2.86)}$ | fish, invertebrates, crustaceans, birds | |
| Oceanic Whitetip Shark | FL = 0 + 0.822 x TL<br>TL= $(KG*1000/0.01000)^{(1/3.07)}$ | fish, invertebrates, crustaceans, birds, reptiles | |
| Pacific Bluefin Tuna | FL = 0 + 0.927 x TL<br>TL= $(KG*1000/0.01549)^{(1/3.02)}$ | fish, invertebrates, crustaceans | brain, eyes, stomach |
| Porbeagle Shark | FL = 0.99 + 0.885 x TL<br>TL= $(KG*1000/0.01047)^{(1/3.03)}$ | fish, invertebrates, sharks | muscles, stomach |
| Sailfish | FL = TL / 1.152<br>TL= $(KG*1000/0.00589)^{(1/3.14)}$ | fish, invertebrates, crustaceans | brain, eyes |
| Salmon Shark | FL = 0 + 0.885 x TL<br>PCL = 0 + 0.878 x FL<br>TL= $(KG*1000/ 0.00933)^{(1/ 3.04)}$ | fish, invertebrates | brain, eyes, muscles, stomach, kidney |
| Shortfin Mako | FL = -1.71 + 0.929 x TL<br>TL= $(KG*1000/0.00646)^{(1/3.03)}$ | fish, invertebrates, crustaceans, sharks | muscle, stomach |
| Skipjack Tuna | FL = (TL - 6.464) / 0.982<br>TL= $(KG*1000/0.01122)^{(1/3.11)}$ | fish, invertebrates, crustaceans | muscle |
| Striped Marlin | FL = TL / 1.083<br>TL= $(KG*1000/0.00550)^{(1/3.15)}$ | fish, invertebrates, crustaceans | brain, eyes |
| Swordfish | FL = TL/1.097<br>TL= $(KG*1000/0.00380)^{(1/3.15)}$ | fish, invertebrates, crustaceans | brain, eyes |
| Whale Shark | FL = (TL - 26.5) / 1.063<br>TL= $(KG*1000/0.00389)^{(1/3.12)}$ | fish, invertebrates, plankton | |
| White Marlin | FL = TL / 1.164<br>TL= $(KG*1000/0.00447)^{(1/3.13)}$ | fish, squids | eyes |
| White Shark | FL = -5.744 + 0.944 x TL<br>TL= $(KG*1000/0.00871)^{(1/3.05)}$ | fish, invertebrates, crustaceans, sharks, birds, mammals | brain, eyes, muscle, stomach |
| Yellowfin Tuna | FL = (TL - 8.831) / 0.967<br>TL= $(KG*1000/0.01479)^{(1/3.01)}$ | fish, invertebrates, crustaceans | |

**Table S4: Parameters of the multilinear regression models for deep and shallow preferred diving depth ($D_{pref}$) of large pelagic predators, day and night.**

| ***Groups (Day)*** | **ΔO2** | **ΔT** | **SST** | **$Z_{eu}$** | **$Z_{meso}$** | **$Log_{10}(Chl)$** | **$Log_{10}(EKE)$** | **$L_{FL}$** | **$r^2$ (p value)** |
|---|---|---|---|---|---|---|---|---|---|
| Shallow diving shark | -0.49 | 3.05 | 3.27 | -0.67 | -0.64 | -65.9 | -23.5 | 0.04 | 0.06 (>0.05) |
| Shallow diving tuna | -0.56 | 2.00 | -1.83 | -1.10 | -0.06 | -72.3 | -24.3 | 0.20 | 0.27 ($p<10^{-2}$) |
| Shallow diving billfish | 0.85 | -6.44 | 3.32 | -0.97 | 0.56 | -44.5 | 27.6 | 0.15 | 0.26 (p>0.05) |
| Deep diving shark | 0.49 | -9.9 | 9.15 | 0.71 | -0.20 | -134 | -5.15 | 0.32 | 0.15 ($<10^{-2}$) |
| Deep diving tuna | **1.26** | -8.73 | **35.4** | -2.77 | **2.46** | **156.9** | 44.9 | **1.70** | 0.73 ($<10^{-24}$) |
| Deep diving billfish | -0.59 | -20.4 | 15.7 | 4.05 | -0.62 | -13.8 | 85.2 | -0.45 | 0.65 ($<10^{-5}$) |

| ***Groups (Night)*** | **ΔO2** | **ΔT** | **SST** | **$Z_{eu}$** | **$Z_{meso}$** | **$Log_{10}(Chl)$** | **$Log_{10}(EKE)$** | **$L_{FL}$** | **$r^2$ (p value)** |
|---|---|---|---|---|---|---|---|---|---|
| Shallow diving shark | 0.09 | 1.27 | 0.50 | -0.56 | 0.23 | -12.1 | -0.63 | 0.03 | 0.44 ($<10^{-3}$) |
| Shallow diving tuna | **-0.52** | -2.83 | 3.56 | 0.80 | **-0.71** | -29.5 | -1.76 | -0.02 | 0.35 ($<10^{-2}$) |
| Shallow diving billfish | 0.26 | -0.66 | 0.59 | -1.15 | 0.43 | -23.4 | -5.59 | -0.03 | 0.04 (>0.05) |
| Deep diving shark | -0.08 | -1.49 | 1.11 | 0.47 | -0.33 | -71.0 | 13.1 | 0.10 | 0.41 ($<10^{-9}$) |
| Deep diving tuna | **-0.36** | **-5.11** | **3.74** | 0.32 | -0.24 | 1.32 | 12.1 | 0.24 | 0.51 ($<10^{-12}$) |
| Deep diving billfish | 0.21 | -1.95 | -1.13 | 0.39 | 0.08 | -7.7 | 11.7 | -0.03 | 0.08 (>0.05) |

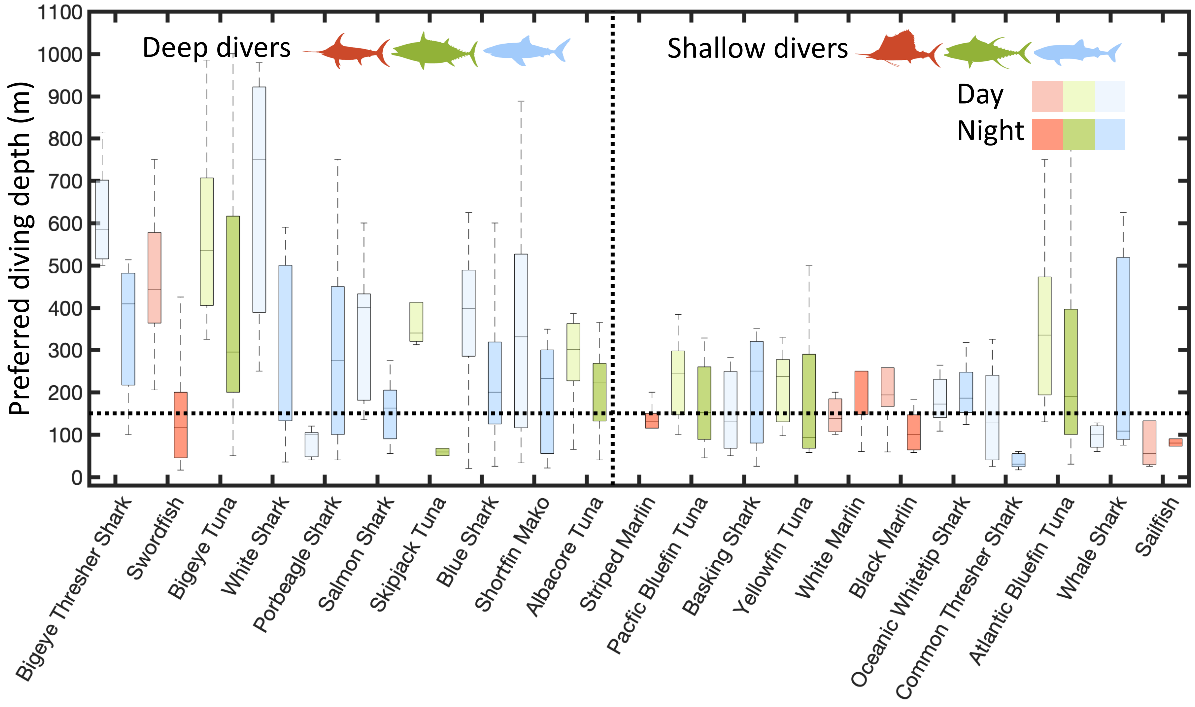


**Figure S1: Daytime and nighttime maximum diving depth ($D_{deep}$) by species and taxonomic group. Results are shown for all data points, omitting three species with less than 3 data points each (blue marlin, spearfish, and southern bluefin tuna). In each box plot, the central mark indicates the median, the top and bottom edges the 75th and 25th percentiles, and the whiskers the full range. Taxonomic groups are shown by different colors (blue=sharks; green=tuna; red=billfish) for daytime (light shades) and nighttime (dark shades).**

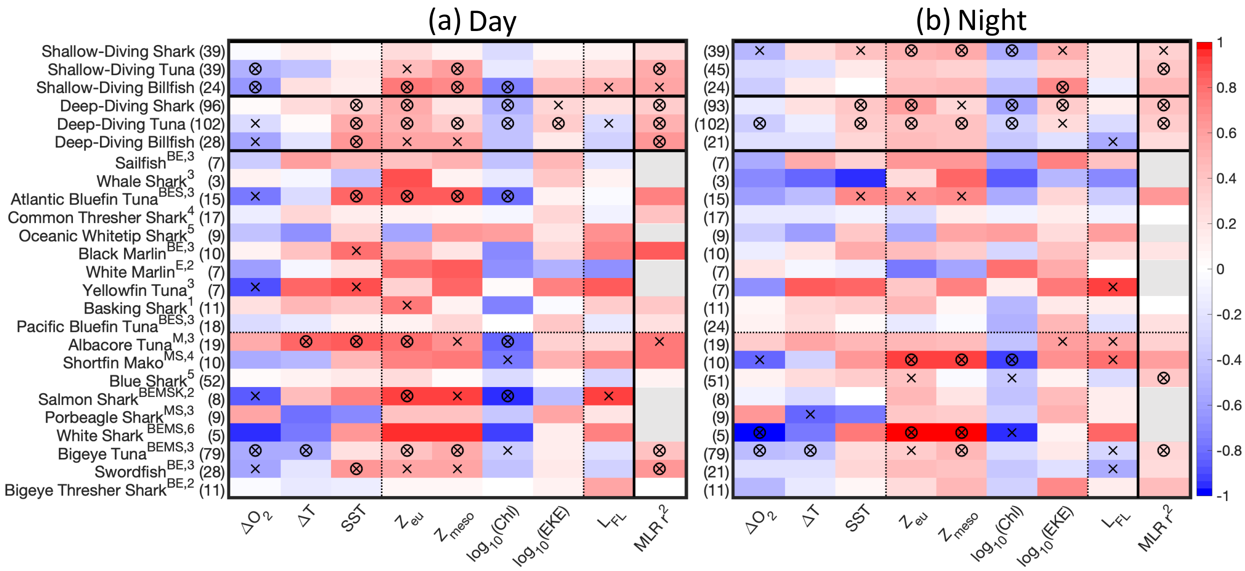


**Figure S2: Correlation between maximum diving depths of large marine predators ($D_{deep}$) and selected environmental drivers for (a) daytime and (b) nighttime. Colors show the Pearson correlation coefficient r between individual drivers and $D_{deep}$ for each category (red=positive correlation blue=negative correlation). In each panel, the leftmost column shows the adjusted $r^2$ of the multilinear regression between all drivers and $D_{deep}$. Gray boxes show cases with too few observations to build a multiple linear regression. Crosses indicate significant correlations at the 1% level, circles at the 0.1% level. The top rows show correlations for aggregated taxonomic groups, separating shallow from deep diving species. The bottom rows show correlations for individual species, ranked from shallowest to deepest. For each species, the letters in superscript indicate presence of thermoregulation apparatuses (B=brain, E=eyes, M=muscles; S=stomach; K=kidney; see also Table S3). The numbers in superscript indicate the level of feeding generalism (i.e., the number of different prey functional groups, see Table S3). Correlations are computed for data ranked A and B in quality, removing outliers and disregarding species with less than 3 samples each (blue marlin, spearfish, and southern bluefin tuna).**

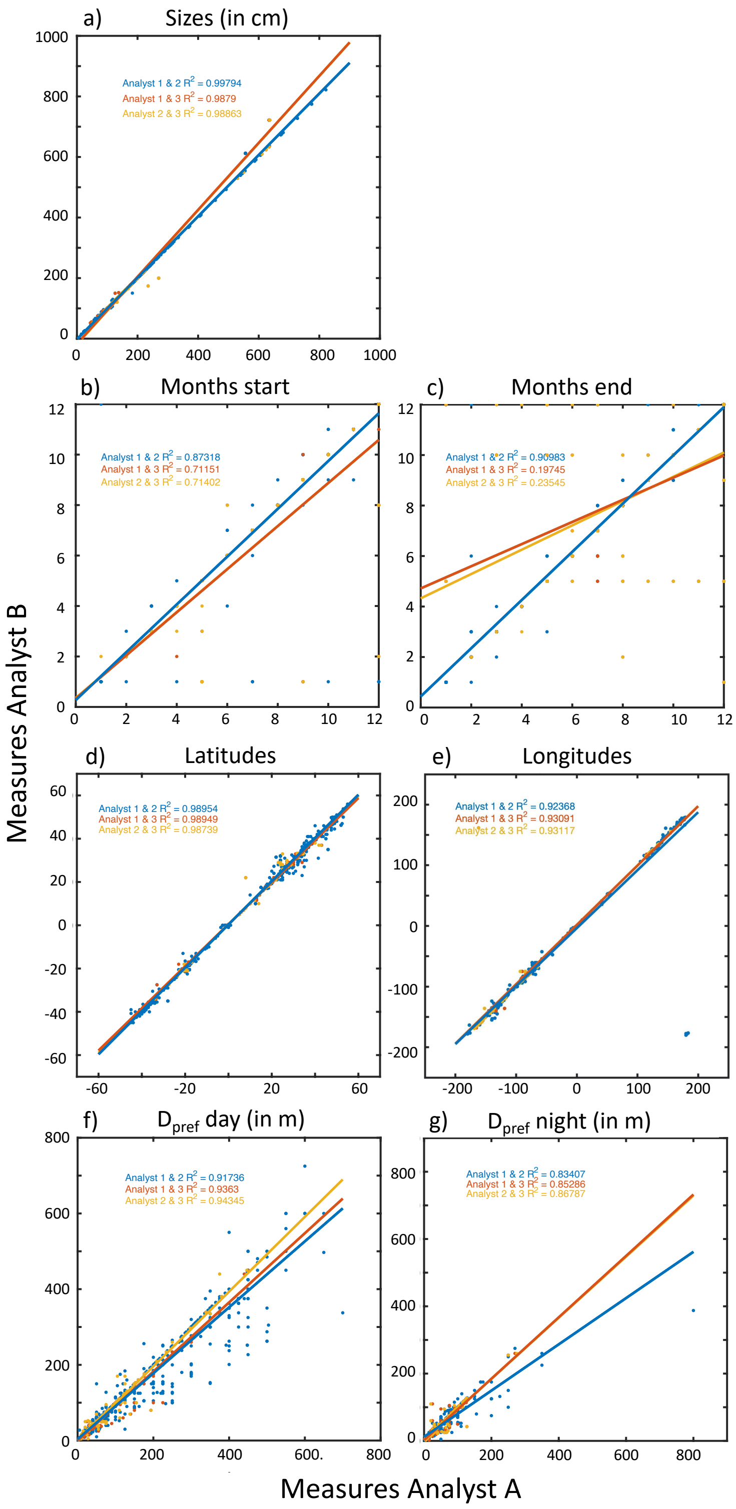


**Figure S3: Comparison of independent data extractions by Analysts 1 to 3: (a) Sample sizes; (b,c) Months range; (d,e) Average latitudes and longitudes; (f,g) Preferred day and night diving depth. Correlation coefficients indicate the match between data extracted by Analyst A (1 or 2) vs. Analyst B (2 or 3).**

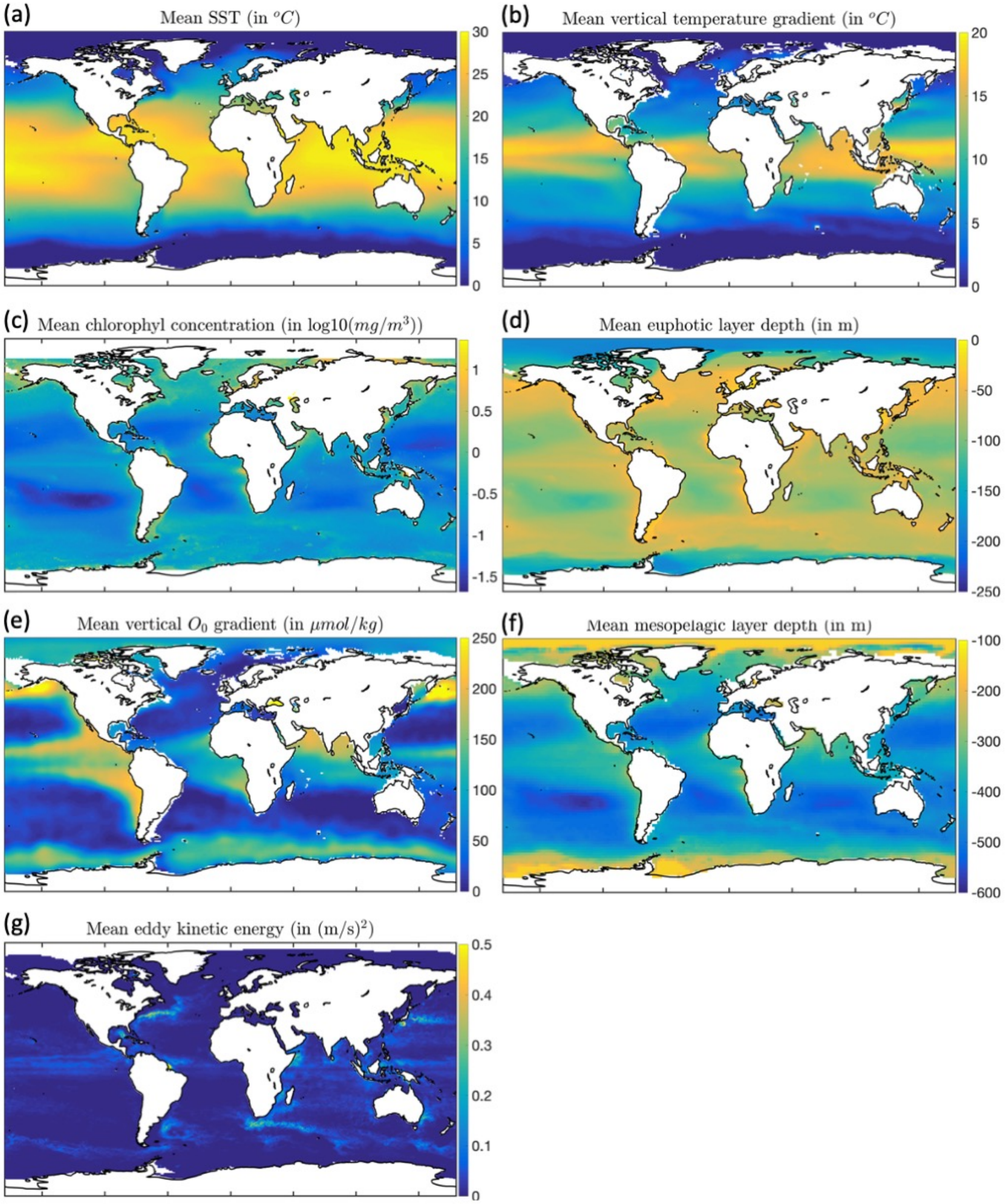


**Figure S4: Mean global distribution of environmental drivers used for comparison with preferred diving depth: (a) sea surface temperature, in °C; (b) temperature gradient between surface and 250m depth, in °C; (c) chlorophyll concentration, in log10(mg/m$^3$); (d) euphotic layer depth, in m; (e) oxygen gradient between surface and 250m depth, in µmol/kg; (f) mesopelagic layer depth, in m; (g) eddy kinetic energy, in (m/s)$^2$.**

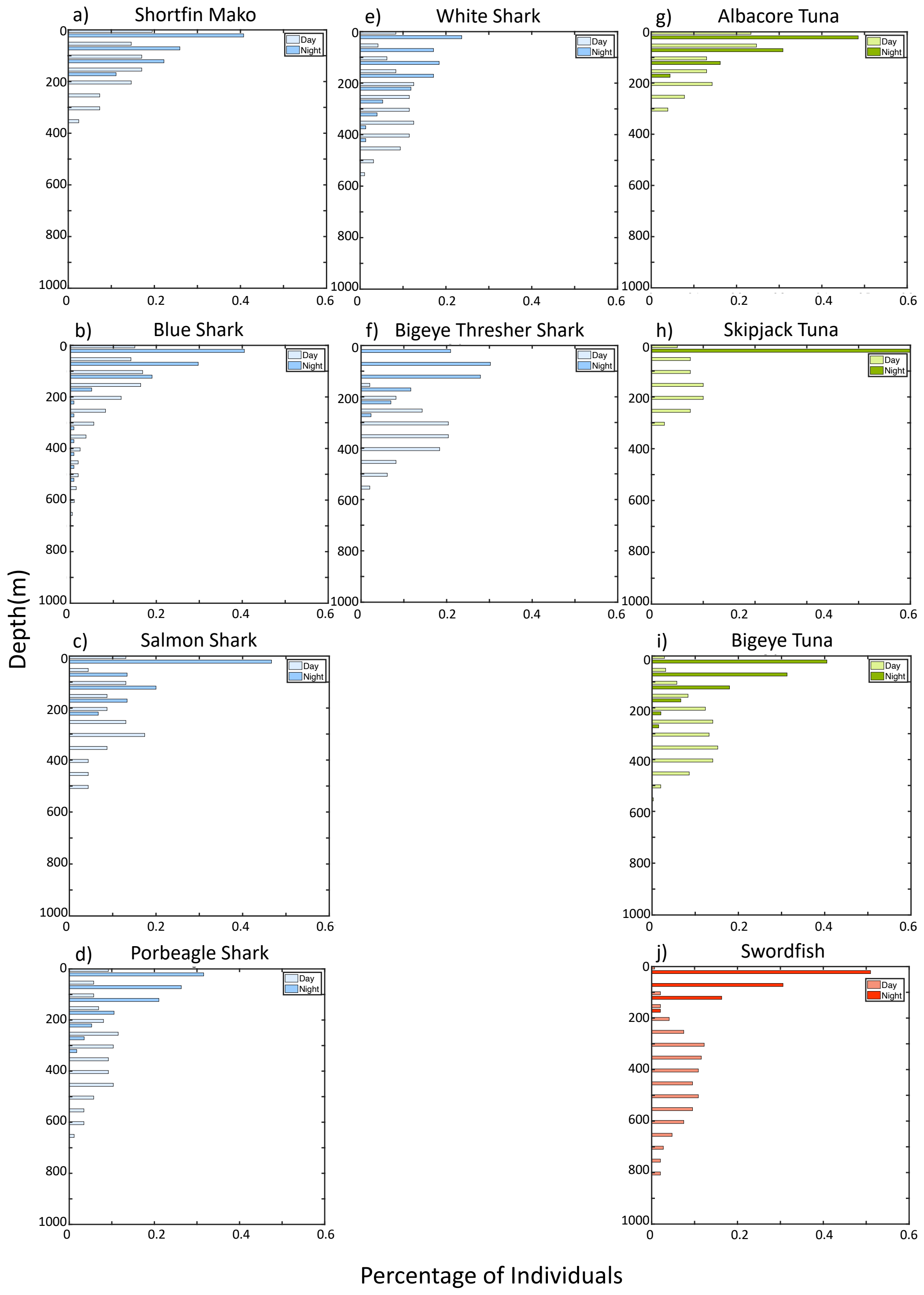


**Figure S5: Distribution of day and night preferred vertical depths ($D_{pref}$) for deep diving large marine predators based on tagging data. Panels (a-f) show shark species (in blue), panels (g-i) show tuna species (in green) and panel (j) swordfish (in red). Dark colors show night depth ranges and light colors day depth ranges.**

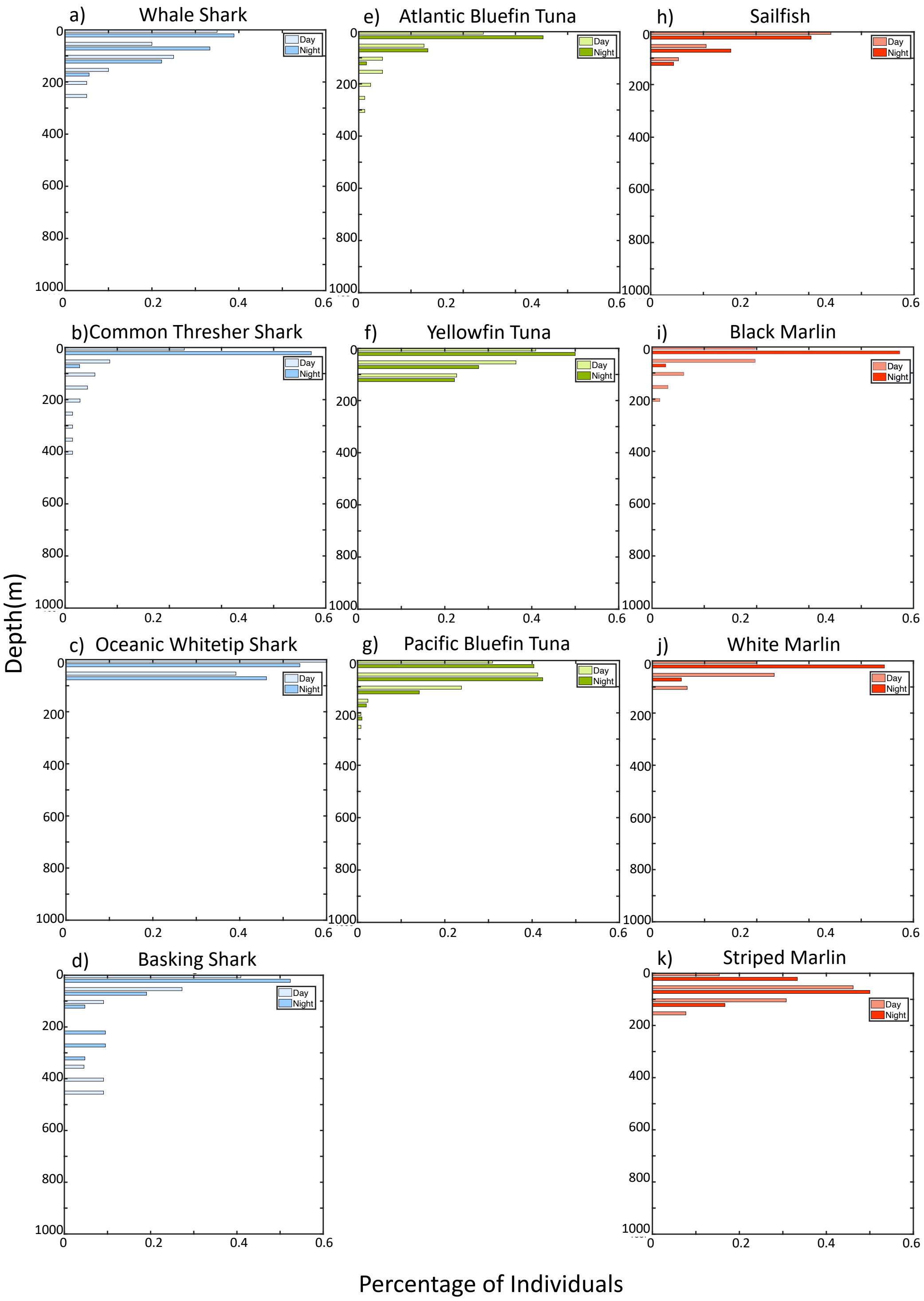


**Figure S6: Distribution of day and night preferred vertical depths ($D_{pref}$) for shallow diving large marine predators based on tagging data. Panels (a-d) show shark species (in blue), panels (e-g) show tuna species (in green) and panels (h-kj) billfish species (in red). Dark colors show night depth ranges and light colors day depth ranges.**

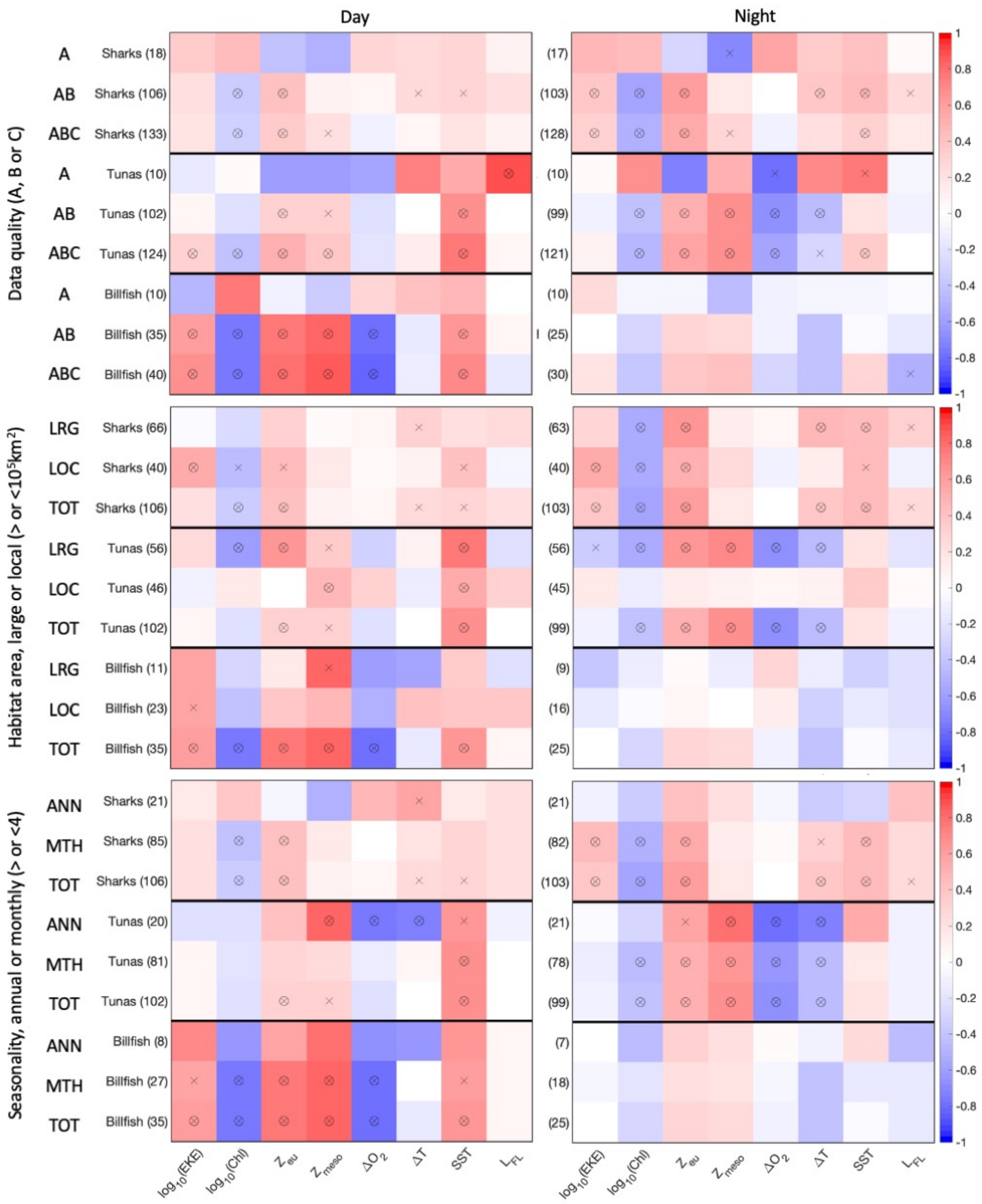


**Figure S7: Sensitivity of the correlation of preferred diving depth for deep divers ($D_{pref}$) with the environment. (a) For data of various quality, including observation from A (best) to C (worst) as ranked by independent analysts. (b) For spatial habitats of various sizes, large (>$10^5$km) or smaller (<$10^5$km), as identified by independent analysts. (c) For various timescales covered by observation, seasons (<=4months) or larger (>4months). For (b) and (c) comparisons include A and B quality observations. Colors indicate the Pearson correlation coefficient r between individual drivers and $D_{pref}$ per species. Crosses indicate 1% significance of correlations, circles 0.1% significance.**

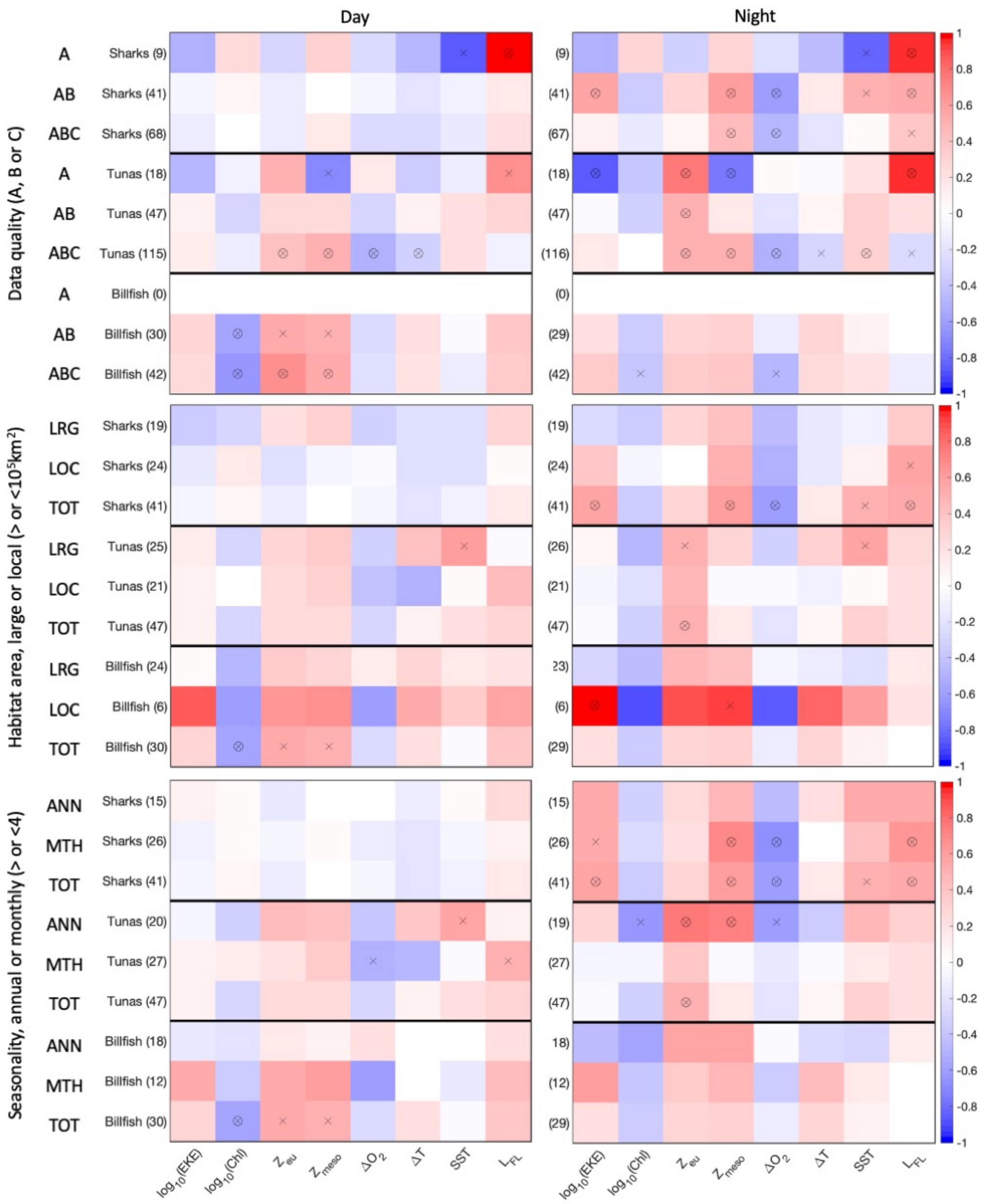


**Figure S8: Sensitivity of the correlation of preferred diving depth for shallow divers ($D_{pref}$) with the environment. (a) For data of various quality, including observation from A (best) to C (worst) as ranked by independent analysts. (b) For spatial habitats of various sizes, large (>$10^5$km) or smaller (<$10^5$km), as identified by independent analysts. (c) For various timescales covered by observation, seasons (<=4months) or larger (>4months). For (b) and (c) comparisons include A and B quality observations. Colors indicate the Pearson correlation coefficient r between individual drivers and $D_{pref}$ per species. Crosses indicate 1% significance of correlations, circles 0.1% significance.**
